# Transcriptomic diversity of cell types across the adult human brain

**DOI:** 10.1101/2022.10.12.511898

**Authors:** Kimberly Siletti, Rebecca Hodge, Alejandro Mossi Albiach, Lijuan Hu, Ka Wai Lee, Peter Lönnerberg, Trygve Bakken, Song-Lin Ding, Michael Clark, Tamara Casper, Nick Dee, Jessica Gloe, C. Dirk Keene, Julie Nyhus, Herman Tung, Anna Marie Yanny, Ernest Arenas, Ed S. Lein, Sten Linnarsson

**Affiliations:** Karolinska Institute, Stockholm, Sweden; Allen Institute for Brain Science, Seattle, Washington; Department of Laboratory Medicine and Pathology, University of Washington, Seattle, Washington, USA

## Abstract

The human brain directs a wide range of complex behaviors ranging from fine motor skills to abstract intelligence and emotion. However, the diversity of cell types that support these skills has not been fully described. Here we used high-throughput single-nucleus RNA sequencing to systematically survey cells across the entire adult human brain in three postmortem donors. We sampled over three million nuclei from approximately 100 dissections across the forebrain, midbrain, and hindbrain. Our analysis identified 461 clusters and 3313 subclusters organized largely according to developmental origins. We found area-specific cortical neurons, as well as an unexpectedly high diversity of midbrain and hindbrain neurons. Astrocytes also exhibited regional diversity at multiple scales, comprising subtypes specific to the telencephalon and to more precise anatomical locations. Oligodendrocyte precursors comprised two distinct major types specific to the telencephalon and to the rest of the brain. Together, these findings demonstrate the unique cellular composition of the telencephalon with respect to all major brain cell types. As the first single-cell transcriptomic census of the entire human brain, we provide a resource for understanding the molecular diversity of the human brain in health and disease.

---

The mammalian brain develops from four compartments—the telencephalon, diencephalon, midbrain, and hindbrain—that give rise to the major structures that underlie the remarkable capabilities of the mature brain. The telencephalon contains the cerebral cortex, hippocampus, and cerebral nuclei and is therefore best studied. Indeed the cortex and hippocampus are widely known as the centers of cognition and memory, respectively, and the cerebral nuclei comprise diverse sets of neuronal clusters that process fear and emotion (the amygdala), as well as movement- and motivation-related information (the basal ganglia). The diencephalon gives rise to the hypothalamus and thalamus, which maintain homeostasis and relay sensory formation, respectively. Posteriorly, the midbrain and hindbrain contain many nuclei that regulate vital functions like cardiac, respiratory, and motor functions. The hindbrain comprises the pons, medulla, and cerebellum, the last of which uniquely emerges from an outgrowth of the developing hindbrain called the rhombic lip. In recent years single-cell sequencing has been used to survey the cell-type diversity present across the entire mouse brain, as well as in select regions and cell types of the human brain, especially the cortex (*1–5*). Nevertheless, cell-type diversity across most of the expansive human brain remains unexplored.

We therefore finely dissected tissue from across the telencephalon, diencephalon, midbrain, and hindbrain and performed single-nucleus RNA sequencing (Fig. 1A; Methods). We set out to sample 100 anatomically distinct locations (hereafter “dissections”) based on the Allen Brain Atlas, each in three donors and in technical duplicates. We ultimately retained 606 high-quality samples covering 106 dissections across ten brain regions, including the most anterior part of the cervical spinal cord in one donor (Fig. S1A; Table S1). We sampled all but twelve dissections from three postmortem donors, although dissections were sometimes combined differently across donors to achieve this goal, e.g., PAG became PAG-DR in one donor. We moreover enriched for neurons using fluorescence-activated cell sorting (FACS) and aimed to collect 90% neurons and 10% non-neuronal cells from each sample. We were not able to achieve this proportion in many non-cortical regions due to an abundance of oligodendrocytes. Total genes captured per cell therefore also varied across dissections partly due to biological variation in cell size; for example, non-neuronal cells are small and yield fewer molecules, e.g., A35r (Fig. S1B). We additionally noted variability across donors, likely reflecting tissue quality (Fig. S1C). We captured a median 6224 cells from each sample and then filtered cells based on their total molecules—as counted by unique molecular identifiers (UMIs)—and percentages of unspliced RNA, as well as doublet scores (Fig. S1D-E).

**Figure 1.**
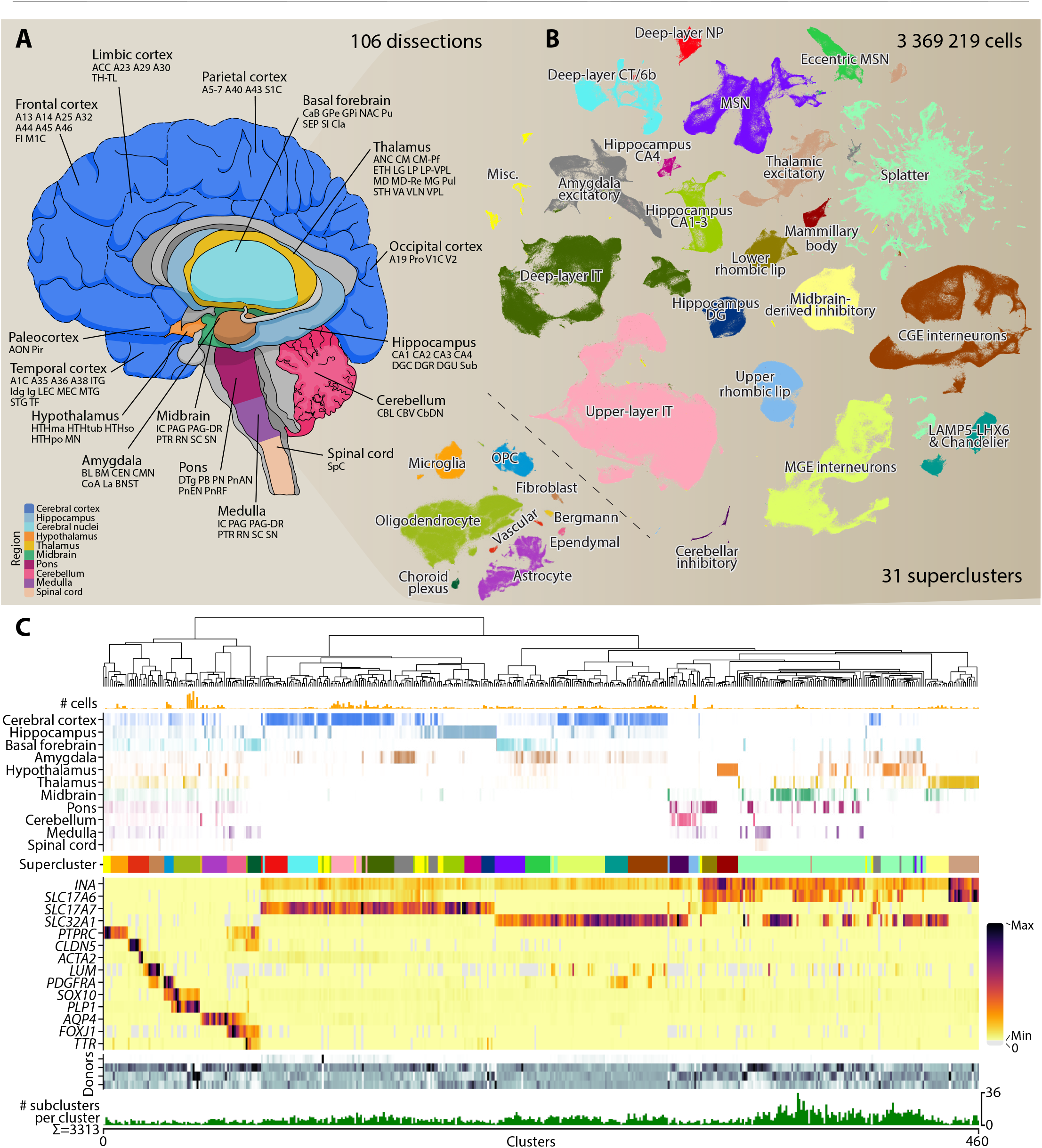
Single-nucleus RNA sequencing reveals transcriptomic diversity across the adult human brain. **(A)** RNA sequencing was performed on 106 dissections replicated in three donors. Schematic shows the location of each dissection across the brain. Abbreviations correspond to the Allen Brain Reference Atlas or cortical Brodmann areas. **(B)** Neurons (right) and non-neuronal cells (inset left) were analyzed separately, and cells on each t-SNE are colored by supercluster. Abbreviations: CA = Cornu Ammonis area; CGE = caudal ganglionic eminence; CT = corticothalamic; DG = dentate gyrus; IT = intratelencephalic; NP = near-projecting; MGE = medial ganglionic eminence; MSN = medium spiny neuron; OPC = oligodendrocyte precursor cell. **(C)** Each supercluster in (B) was analyzed separately, yielding the 461 clusters arranged in the dendrogram. Attributes are plotted for each cluster from top to bottom: bars represent the number of cells; color depth represents the proportion of cells that derive from each brain region; color supercluster is colored as in (B); heatmap shows expression levels for genes that mark neurons (INA), glutamatergic (SLC17A6, SLC17A7) and GABAergic (SLC32A1) neurons, immune cells (PTPRC), vascular cells (CLDN5, ACTA2), fibroblasts (LUM), OPCs (PDGFRA, SOX10), oligodendrocytes (SOX10, PLP1), astrocytes (AQP4), ependymal cells (FOXJ1), and choroid plexus (FOXJ1, TTR); color depth represents the proportion of cells that derive from each donor; bars represent the number of subclusters produced by analyzing the cluster separately.

We then pooled high-quality cells from all dissections into a single dataset for graph-based clustering (Methods). Harmony was used to integrate cells across donors (*6*). The resulting clusters were split into two datasets based on the dendrogram that largely corresponded to neurons and non-neuronal cells, and each dataset was re-clustered. We used the Paris algorithm to build a dendrogram from these clusters (*7*), and then cut the dendrogram to identify “superclusters” that corresponded to islands on a two-dimensional representation calculated by t-distributed stochastic neighbor embedding (t-SNE, Fig. 1B). Neurons and non-neuronal cells were divided into 21 and 10 superclusters, respectively. Each was manually annotated based on the literature and their regional composition (Table S2) with one exception, a supercluster that contained neurons from most brain regions. In the tradition of naming neurons for their shapes, we named them *splatter neurons* based on their appearance on the two-dimensional embedding. Each supercluster was separately reanalyzed, resulting in a final set of 461 clusters. A dendrogram of these clusters mirrored previous work in the mouse: neurons split separately from non-neuronal cells and into two clades (Fig. 1C) (*1*). One clade contained telencephalic excitatory neurons that express glutamatergic genes like SLC17A6 and SLC17A7. The other contained telencephalic inhibitory neurons that express GABAergic genes like SLC32A1, as well as all excitatory and inhibitory diencephalic, midbrain, and hindbrain neurons. We additionally analyzed each cluster separately to produce a more detailed but less robust clustering that contained 3313 clusters, which we subsequently refer to as “subclusters” (Fig. 1C).

The distributions of superclusters across dissections exposed three experimental caveats of our work (Fig. 2A). First, dissections were not always replicable across donors, partly because their anatomical borders were challenging to distinguish. For example, although telencephalic superclusters contained some hypothalamus and thalamus cells, we suspect that these cells derived from telencephalic tissue bordering these regions. Second, we sequenced multiple dissections together at once, and sequencing reads were occasionally misassigned across samples (*8*). Although we implemented a preprocessing pipeline that computationally filtered these “index-hopping” reads (Methods), the filter was likely imperfect. Some cells from the globus pallidus (GPi) clustered with hindbrain neurons (Fig. 2A) but contained fewer UMIs than similar cells. These cells also derived from a sample sequenced together with a hindbrain dissection, implying read misassignment. Third, our computational approach to determine superclusters left some rare cell types poorly connected on the graph. These cells were dissimilar to one another but ended up in a single supercluster called *miscellaneous*. For example, this supercluster contained lymphocytes that were distinct from other cell types in the brain.

**Figure 2.**
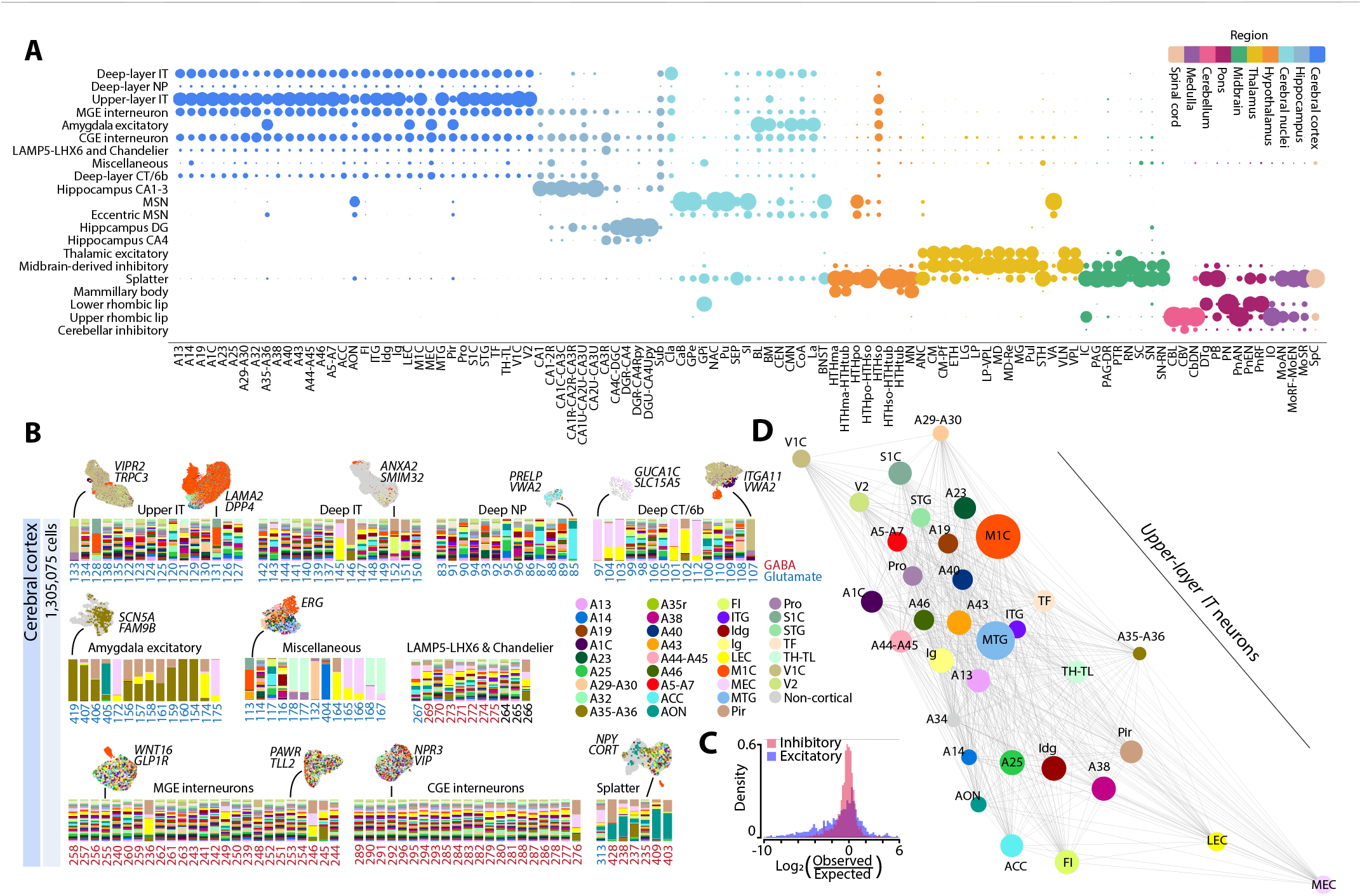
Superclusters and clusters are differentially distributed across brain regions. (A) Dot plot represents supercluster distributions across dissections. Within each column, dot size is proportional to the percentage of cells from the indicated dissection that belong to each supercluster. Dot color indicates region according to the legend. (**B**) Stacked bar plots show the relative contributions of neurons from each cortical dissection, colored according to the legend. Only clusters larger than 100 cells with at least 1% contribution from the cerebral cortex are shown. A Uniform Manifold Approximation and Projection (UMAP) embedding is also shown for select clusters, where non-cortical cells are colored grey. Clusters are labeled with two of their most enriched genes. Red labels indicate clusters expressing inhibitory GABAergic vesicular transporter *SLC32A1*; blue clusters express excitatory glutamatergic vesicular transporter *SLC17A6* or *SLC17A7*. (**C**) Histograms show the observed number of cells divided by the expected number of cells (given the number of cells sampled overall and for each region), for each cluster and dissection. (**D**) Neighbor graph represents *upper-layer IT* cells from all cortical dissections. Node size is proportional to the number of cells. The weight of an edge between two dissections is proportional to the percentage of cells in the dissections that are neighbors.

Nevertheless, supercluster distributions revealed large-scale similarities among adult neurons that reflected their developmental history (Fig. 2A). Superclusters were largely exclusive to either telencephalic or non-telencephalic dissections, and some were additionally specific to regions like the hippocampus, cerebral nuclei, thalamus, or hypothalamus. Additionally, the presence of three superclusters in multiple regions reflected migration during development. First, cortical interneurons migrate from the medial and caudal ganglionic eminences (MGE and CGE, respectively) during development. Correspondingly, *MGE* and *CGE interneurons* distributed across the telencephalon. Our data suggested that these cells migrate more widely than appreciated; hypothalamus, thalamus, and midbrain contained cells from this supercluster. This observation might reflect imprecise dissection or transcriptomic convergence during development; some of the same transcription factors like LHX6 generate inhibitory neurons in the hypothalamus (*9*). However, neuronal migration from the ganglionic eminences into the thalamus occurs specifically in human tissue, raising the possibility that some of these cells have indeed migrated (*10*). Second, a supercluster of inhibitory neurons was distributed mainly in the thalamus and midbrain, likely corresponding to the SOX14+ neurons that migrate from the embryonic midbrain to populate the thalamus (*11*). Our data suggested that some of these cells also migrate to the hypothalamus and pons; accordingly, we localized marker genes for these cells in the pons with RNAScope *in situ* hybridization (Fig. S2). Third, two superclusters appeared to derive from the rhombic lip, which produces not only the cerebellum but also neurons that migrate to specific nuclei in the pons and medulla (*12*). One of these superclusters contained cerebellar granule neurons that derive from the upper rhombic lip. The second contained neurons from the pontine nucleus that derive from the lower rhombic lip.

We used supercluster and cluster labels to explore the cellular composition of each brain region. Most of the cortex is organized in six layers that contain diverse interneurons and excitatory neurons that project within and outside the telencephalon. Most cortical cells were accordingly MGE- and CGE-derived interneurons or layer-specific excitatory neurons: intratelencephalic (*upper-layer* and *deep-layer IT*), near-projecting (*deep-layer NP*), and corticothalamic (*deep-layer CT/6b*) (Fig. 2A). Although cortical layers two through four are commonly considered “upper” and layers five and six “deep,” both *upper-layer IT* and *deep-layer IT* superclusters contained putative Layer-4 clusters (Fig. 2B; #133, 138, 141). Layer-5 extratelencephalic neurons (ET) were found in the *miscellaneous* supercluster (Fig. 2B; #113-114, 116-118). One Layer-5 ET cluster—found mostly in frontal insula and anterior cingulate cortex—expressed von Economo markers like *BMP3* and *ITGA4* (Fig. 2B; #117) (*13*). Although we also observed *splatter* neurons in the cortex, most derived from the anterior olfactory nucleus (AON), a dissection that likely contained some basal-ganglia tissue. Only one *splatter* cluster contained cells from across the cortex and might represent *SST*-*CHODL* long-range projecting neurons, expressing SST, NPY, and high levels of NOS1 (Fig. 2B; #235) (*14*).

Strikingly, layer-specific excitatory neurons varied more across dissections than interneurons (Fig. 2C). Most interneuron clusters contained cells from all cortical dissections, and the dissections were nearly indistinguishable on a Uniform Manifold Approximation and Projection (UMAP) embedding of these clusters. In contrast, some excitatory clusters were enriched in specific dissections, such as visual cortex (V1C), Brodmann areas 5-7 (A5-A7), frontal insula (FI), anterior cingulate cortex (ACC), or entorhinal cortex (A35-A36, MEC, and LEC). These specializations occurred in different superclusters: whereas ACC cells were enriched in one *deep-layer NP* cluster (#85), A5-A7 cells were enriched in two *upper-layer IT* and one *miscellaneous* Layer-5 ET cluster (#128, 131, 113). V1C exhibited distinctive *deep-layer CT/6b* and *upper-layer IT* clusters (#107, 133), the latter of which we annotated as V1C’s distinctive *TRPC3*+ Layer 4 (*15*). Entorhinal and piriform cortex (Pir) not only contained distinctive *deep-layer IT* and *deep-layer CT/6b* clusters, but also neurons that clustered with *amygdala excitatory* cells (Fig. 2B). These regions also contained distinct interneuron clusters (Fig. 2B; #236, 246, 276). Notably, entorhinal and piriform cortex is considered part of the more evolutionarily conserved allocortex that contains only three cellular layers. To more broadly assess transcriptomic variation across different cortical dissections, we visualized cortical *upper-layer IT* neurons as a graph where each node represented a dissection, and each edge represented the percentage of nearest neighbors shared between two dissections (Fig. 2D). Cortical dissections ranged transcriptomically between entorhinal and visual cortex, and dissections from nearby anatomical locations often neighbored one another, revealing a transcriptional gradient across the cortex.

Other telencephalic regions exhibited both cortex-related and region-specific superclusters. For example, the hippocampus contained *MGE* and *CGE interneurons* in addition to three superclusters related to its anatomical subdivisions: dentate gyrus (*DG*) and *cornu ammonis* subfields 1-4 (*CA1-3, CA4*) (Fig. 2A; Fig. S3A). We also observed some layer-specific excitatory neurons that mostly derived from the cortex-adjacent subiculum. The cerebral nuclei were more heterogeneous in accordance with their diverse neuronal circuitry. Claustrum contained mostly cortical superclusters, whereas the basal ganglia— caudate, putamen, nucleus accumbens, and the external and internal segments of the globus pallidus (CaB, Pu, NAc, GPe, Gpi)—contained mostly medium spiny neurons (*MSNs*, Fig. 2A). Interestingly, we observed two *MSN* superclusters, one of which we putatively annotated as the recently described *CASZ1*+ “eccentric” MSNs (Fig. S3B-C) (*5*). *MSN* clusters were relatively specific to— although differentially distributed across (Fig. S3A)—the basal ganglia, whereas *eccentric MSNs* were more evenly distributed throughout the cerebral nuclei (Fig. 2A). Most *eccentric MSNs* expressed only *DRD1*, but many in the basal ganglia co-expressed *DRD1* and *DRD2*, suggesting regional specialization of a more broadly distributed cell type (Fig. S3D-E). In addition to *MSNs*, the amygdala contained cortical superclusters—interneurons and some layer-specific excitatory neurons—and its own excitatory supercluster. Both *MSNs* and *amygdala excitatory* neurons were differentially distributed across amygdala dissections, implying that individual nuclei exhibit unique neuronal compositions (Fig. S3A). The bed nucleus of the stria terminalis (BNST), considered extended amygdala, appeared particularly distinct and contained only *MSNs* and *splatter* neurons. Indeed most amygdaloid *splatter* neurons derived from BNST.

Outside the telencephalon, the thalamus contained three major superclusters: a region-specific *thalamic excitatory* supercluster, *midbrain-derived inhibitory* neurons, and *splatter* neurons, as well as the cortical interneurons detailed earlier (Fig. S3A). Similar to the amygdala, *thalamic excitatory* cells varied more across dissections than *midbrain-derived inhibitory* clusters which contained cells from all thalamic dissections. We also observed *MSNs, amygdala excitatory* neurons, and *hippocampal DG* neurons in the thalamus that we attributed to dissection imprecision. Among all regions profiled, the cerebellum was least complex; although we observed some granular-, molecular-, and purkinje-layer interneurons, our samples were dominated by the highly abundant *upper rhombic lip* granule neurons (Fig. 2A). Finally, the hypothalamus, midbrain, and other hindbrain regions contained some abundant, relatively homogeneous neurons—*mammillary* neurons, *midbrain*-*derived inhibitory* neurons, and rhombic lip-derived neurons, respectively—but most other neurons in these regions were *splatter* neurons.

*Splatter* neurons represented a strikingly heterogenous group of cells that did not share a highly specific, broadly expressed gene module. 291,883 cells were distributed over 92 clusters, although many cells fell into several large clusters centrally located on the embedding that expressed no specifically enriched genes (Fig. 3A). Except for a subset of hypothalamic cells, most *splatter* neurons did not differ from other neurons in terms of quality metrics like total UMI counts or unspliced RNA fractions (Fig. S4A-B). The *splatter* supercluster uniquely comprised both inhibitory and excitatory cells: 22.1% of cells expressed the GABA transporter *SLC32A1*, 39.3% of cells expressed the predominant glutamatergic transporter *SLC17A6*, and 1.4% of cells co-expressed these two genes (Fig. 3B; Fig. S4C). The dataset moreover included cholinergic (*SLC5A7*+) and glycinergic (*SLC6A5*+) neurons, as well as more than half of all hypothalamic neurons, many of which expressed low levels of the hormone vasopressin (*AVP*, Fig. 3C; Fig. S4D-E). A gene ontology analysis of the most variable genes suggested that these cells were differentiated in particular by extracellular matrix- and synapse-related genes (Fig. S4F), reflecting the diversity of regions observed and neurotransmitters expressed in this supercluster.

**Figure 3.**
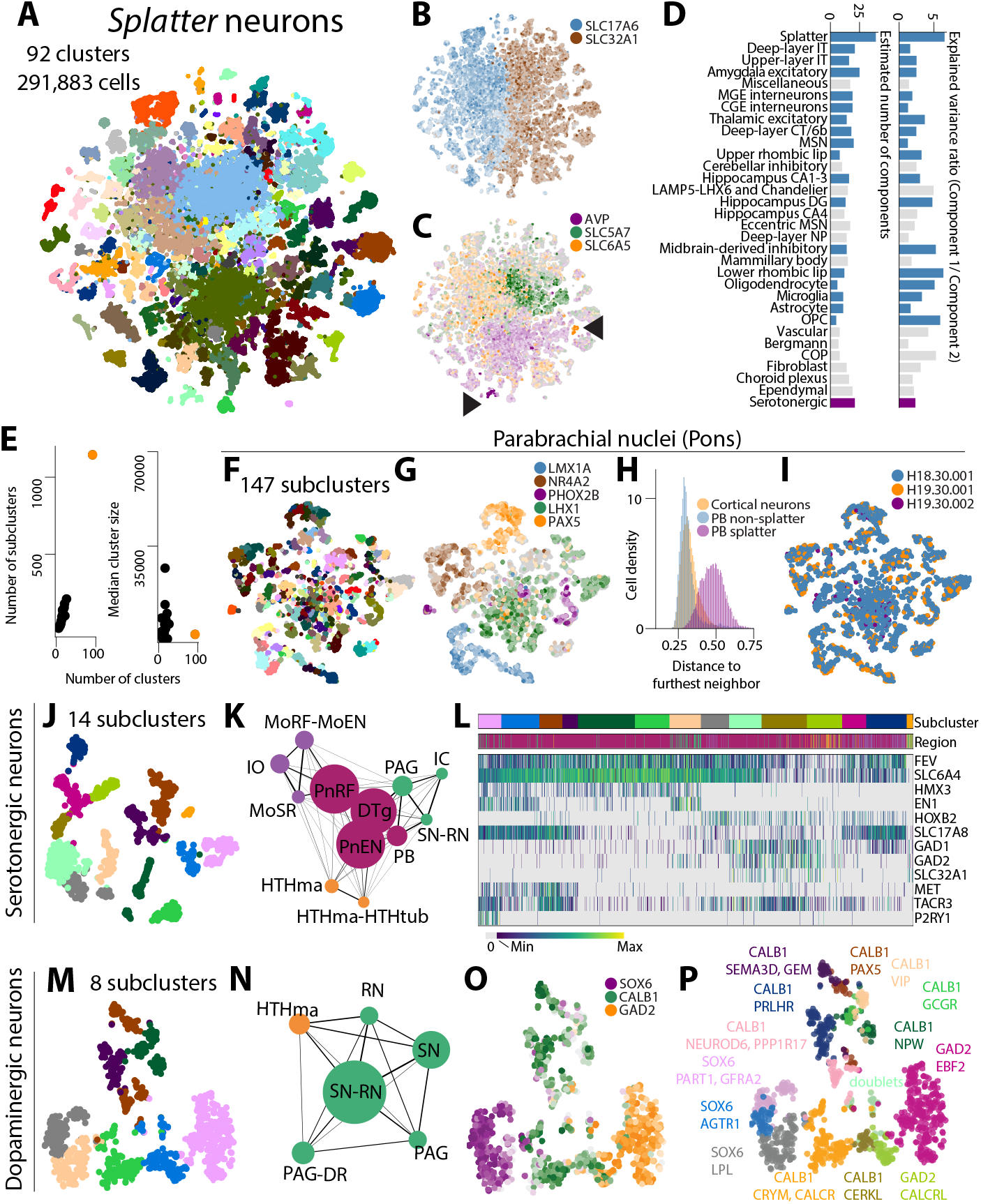
Splatter neurons are highly complex and likely undersampled. (**A**) *Splatter* neurons are colored on the t-SNE by cluster. (**B-C**) Cells are colored by expression levels of the gene they express most highly among those in the legend. Darker color indicates higher expression; gray indicates no expression. **(C)** Arrows indicate clusters with high AVP and SLC6A5 expression, likely true vasopressin-releasing and cholinergic clusters, respectively. (**D**) For each supercluster, the bar represents the optimal number of principal components to describe the dataset (left) or the ratio between the fractions of variance that the first two principal components explain (right). Blue bars indicate superclusters downsampled to 50,000 cells for analysis; grey superclusters contain fewer than 50,000 cells. Purple bar denotes a subset of *splatter* cells, serotonergic neurons. (**E**) For each supercluster, total clusters is plotted against total subclusters (left) or mean cluster size (right). *Splatter* is orange. (**F**) *Splatter* cells from the parabrachial nuclei are colored by cluster on the t-SNE. (**G**) Same as (B). (**H**) For cortical neurons, parabrachial *splatter* neurons, and all other parabrachial neurons, histograms summarize the distributions of Euclidean distances between cells and their 25^th^ nearest neighbors. Distances are normalized by the dataset’s maximum distance. (**I**) Cells are colored by donor. (**J**) Serotonergic neurons are colored by cluster on the t-SNE. (**K**) A graph represents the dissections that constitute 95% of the serotonergic neurons shown in (J). Node size is proportional to the number of cells from each dissection. The weight of an edge between two dissections is proportional to the percentage of cells in the dissections that are neighbors. (**L**) A heatmap represents expression levels across serotonergic cells for select genes. Top bar indicates the subcluster for each cell, colored as in panel (J). The second bar indicates the region of origin for each cell, colored as in panel (K). (**M**) Dopaminergic neurons are colored by subcluster on the t-SNE. (**N**) Same as (K) but for dopaminergic neurons. (**O**) Same as (B). (**P**) Subclusters in (M) were split and further reanalyzed to produce finer subtypes. Cells are colored by subtype.

To assess the complexity of these cells relative to other superclusters, we estimated the number of principal components necessary to represent each supercluster by performing PCA on permuted datasets. We then compared the variance explained by each component in the original and permuted datasets (Fig. S4G; Methods). Nearly twice as many principal components were necessary to capture *splatter* neurons than other superclusters (Fig. 3D). However, six times the variance was explained by the first principal component than the second, a number more similar to homogeneous superclusters like *upper rhombic lip, oligodendrocytes*, and *OPCs* (Fig. 3D). *Splatter* cells therefore exhibit highly variable but relatively uncorrelated expression. To better capture this heterogeneity, we performed all further analysis using the subcluster labels for these cells (Fig. S4H). Although the median number of cells in each *splatter* cluster was smaller than for many other neuronal superclusters, the 92 clusters yielded 1145 subclusters, over 900 more than any other supercluster (Fig. 3E). In agreement with our findings across the whole brain, *splatter* neurons were most similar across neighboring anatomical and developmental compartments, reflected both by the proportions of subclusters shared across regions and the distributions of cells from each region across the t-SNE (Fig. S4I-J).

*Splatter* neurons were nevertheless highly complex even within individual dissections. We investigated these neurons in the parabrachial nuclei, where we sampled a particularly high proportion of *splatter* neurons. The parabrachial nuclei are located in the pons and form a relay center for interoceptive and exteroceptive sensory inputs like taste, respiration, and thermoregulation. In addition to glia, our parabrachial dissections contained cells from five neuronal superclusters consistent with their position bordering the midbrain: *midbrain-derived inhibitory, upper rhombic lip, lower rhombic lip, cerebellar inhibitory*, and *splatter* (Fig. S4K-N). We found 147 *splatter* subclusters with more than five cells in the parabrachial nuclei (Fig. 3F; Fig. S4O). Our analysis captured three populations known to be spatially restricted within the murine parabrachial nuclei: *FOXP2*+ neurons, *LMX1B*+ neurons, and *PAX5*+ neurons (*16, 17*). The former two populations are more specifically labeled in our dataset by *NR4A2* and *LMX1A* (Fig. 3G). *LMX1A*+ cells include the two previously described *CALCA*+ and *SATB2*+ positive populations (Fig. S4P). *PHOX2B* moreover identifies populations ventral and medial to the parabrachial nuclei in mice (Fig. 3G) (*16*). Our observations therefore suggest broad conservation of parabrachial composition across mice and humans.

However, we observed more GABAergic neurons than expected in the parabrachial nuclei, which contain primarily glutamatergic neurons (Fig. S4Q). Most GABAergic neurons belonged to a centrally located cluster that broadly expressed *LHX1* (Fig. 3G). Suspecting that these neurons originated from neighboring nuclei, we used our complete dataset to identify the most common dissection for each *splatter* subcluster that contained more than 20 parabrachial cells. Although 78 clusters contained primarily parabrachial cells, seven regions predominated the remaining clusters (Fig. S4R-S). Two predominantly hypothalamic clusters contained few UMIs and were therefore likely debris (Fig. S4O,S). Many *LHX1*+ cells belonged to clusters populated by cells from the dorsal tegmental nucleus (DTg), a region of the brain involved in spatial navigation which contains many GABAergic cells. The *LHX1*+ neurons might then represent regions bordering the parabrachial nuclei like the lateral DTg. These observations emphasize that neuronal diversity varies considerably even across neighboring nuclei.

Our analysis of parabrachial *splatter* neurons suggests three broader properties of the highly heterogeneous *splatter* supercluster. First, these neurons were more heterogeneous than other superclusters sampled in the same dissection. *Splatter* neurons in the parabrachial nuclei exhibited greater distances from their 25^th^ nearest neighbors than did neurons from other superclusters, whose distances were more like those between cortical neurons (Fig. 3H; Fig. S4N). Secondly, *splatter* neurons might not reflect their developmental origins, unlike other superclusters. Many *FOXP2*+ parabrachial neurons derive from the rhombic lip (*16*), but we found that most *FOXP2+* neurons clustered with *splatter* neurons—rather than with one of the rhombic-lip derived superclusters—implying that some *splatter* neurons are also rhombic-lip derived. Our parabrachial dissection also contained *FOXP2*+ *lower rhombic lip* neurons, but these cells were likely too few in number to represent all rhombic lip-lineage neurons in the parabrachial nuclei. Lastly, our analysis suggested that the immense complexity of *splatter* neurons reflects highly reproducible cell types. After recalculating the t-SNE without batch correction, the embedding demonstrated striking integration across donors, particularly across the donors from which most parabrachial neurons were sampled (Fig. 3I). The fine structure of *splatter* neurons was therefore highly reproducible between donors and likely reflects true cell-type specialization.

Because many *splatter* clusters were defined by their neurotransmitter expression, we also investigated subcluster diversity by neurotransmitter type. Serotonergic neurons are found across the brain stem and generally express the transcription factor *FEV* and the serotonin reuptake transporter *SLC6A4*. In our analysis, *FEV* and *SLC6A4* were highly expressed in cluster 397, which comprised 14 subclusters that derived primarily from the pons, midbrain, medulla, and hypothalamus (Fig. 3J-K; Fig. S5A-I). All cells expressed *FEV* except a small cluster of 20 *PAX6*+ cells and some *PITX2*+ hypothalamus cells (Fig. 3L; Fig. S5I). As previously described in mice, all subclusters could be grouped into rostral-(*HMX3*+) and caudal- (*HOX*+) derived populations, and rostral neurons could be further split into *EN1*+ and *EN1*-populations (Fig. 3L) (*18*). Similar to a recent single-cell study of mouse serotonergic neurons in the midbrain raphe nuclei (*19*), our analysis revealed combinatorial expression of glutamatergic and GABAergic genes *SLC17A8, GAD1*, and *GAD2* in addition to neuropeptides like *PENK* and *TRH* (Fig. 3L; Fig. S5G-I). We also similarly observed *FEV*+ cells with little *SLC6A4* expression, which the authors speculate might indicate neurotransmitter plasticity. However, we additionally observed a substantial *HMX3*+ population that expressed neither *SLC17A8, GAD1*, nor *GAD2*, as well as a population that expressed the GABAergic transporter *SLC32A1*, suggesting co-release of *GABA* and serotonin. These observations might represent either region- or species-specific specializations. Notably, one subcluster in our analysis appeared to correspond to the distinct *P2RY1*+*MET*+*TACR3*+ cluster located near the cerebral aqueduct in mice (*19*). This subcluster contains mostly cells from the periaqueductal gray, suggesting this cell type and its unique electrophysiological and projection properties are conserved in humans.

We also further investigated *splatter* cluster 395 that was enriched in dopaminergic markers *TH, SLC6A3*, and *DDC* and comprised eight subclusters containing mostly midbrain—substantia nigra, red nucleus, and periaqueductal grey—and a few hypothalamic neurons(Fig. 3M-N; Fig. S5J-N). *SOX6, CALB1* and *GAD2* expression defined three major classes of these neurons, a subset of which we reanalyzed and further divided into 14 subtypes (Fig. 3O-P; Fig. S5O; Methods). We identified one small subtype as doublets based on oligodendrocyte-marker expression, highlighting the challenges of performing quality control within this complex supercluster. The remaining subtypes revealed greater molecular complexity than previously reported by Kamath et al., who sorted *NR4A2*+ nuclei from the substantia nigra and found ten subtypes and two classes (*SOX6+* and *CALB1+)* of dopaminergic neurons (*4*). Using a classifier trained on Kamath’s data, we found that seven of our 14 *TH*+ subtypes corresponded to eight previously defined subtypes (Fig. S5P-S; Methods). Our broader sampling allowed the detection of seven new *TH*+ subtypes including one in the substantia nigra (*SOX6, LPL)* and four *CALB1*+ subtypes (*PAX5, NPW, GCCR, VIP*) in the periaqueductal grey and ventral tegmental area surrounding the red nucleus (Fig. 3P; Fig. S5O). Similar to serotonergic neurons, *CALB1* subtypes in the periaqueductal grey expressed different combinations of neuropeptides like *VIP, NPPC, NPW* and *PENK* (Fig. S5T). Finally, we found two *GAD2*+ clusters (*CALCR* and *EBF2*) that defined the molecular signature of a third class of human *TH*+ neurons that co-express GABAergic markers (Fig. S5O,U) and can release GABA (*20*). Our results are thus consistent with midbrain TH+ neurons exhibiting a range of transcriptional profiles from dopaminergic to GABAergic phenotypes.

To explore the extensive cellular heterogeneity we observed in the midbrain and hindbrain, we used spatial transcriptomics to characterize gene-expression distributions across three tissue sections A, B, and C (Fig. 4; Fig. S6). Enhanced ELectric (EEL) fluorescence in-situ hybridization (FISH) enabled us to profile 445 genes in each section and thereby define anatomical locations based on their cellular composition (*15*). Dopaminergic cells demarcated the parabrachial pigmented nucleus (PBP, Fig. 4A: A1), substantia nigra (SN, Fig. 4A: A2,A3), and ventral tegmental area (VTA, Fig. 4B: B1,B2). GABAergic cells within the substantia nigra further distinguished *pars compacta* and *pars reticulata*; the former lacked GABAergic cells. Glutamatergic *PVALB*+ cells, molecularly distinct from the GABAergic *PVALB*+ neurons scattered throughout the VTA, occupied the red and pontine nuclei (Fig. 4A-B: B3,B13; Fig. S6A) (*21*). Additionally, noradrenergic cells marked the *locus coeruleus* (Fig. 4C: C2). Although we profiled some of these locations with single-nucleus RNA sequencing, gene detection was too sparse to systematically localize our clusters. However, we identified two spatial populations that matched RNA-seq clusters. First, a dense cluster of glutamatergic *PVALB*+ neurons in the pontine nucleus also expressed *HOPX* and therefore corresponded to *lower rhombic lip* neurons, as expected (Fig. S6C). Secondly, *CALB2*+ dopaminergic cells in the parabrachial pigmented nucleus co-expressed dopaminergic and GABAergic markers, similar to a dopaminergic subtype we sequenced (Fig. S6A).

**Figure 4.**
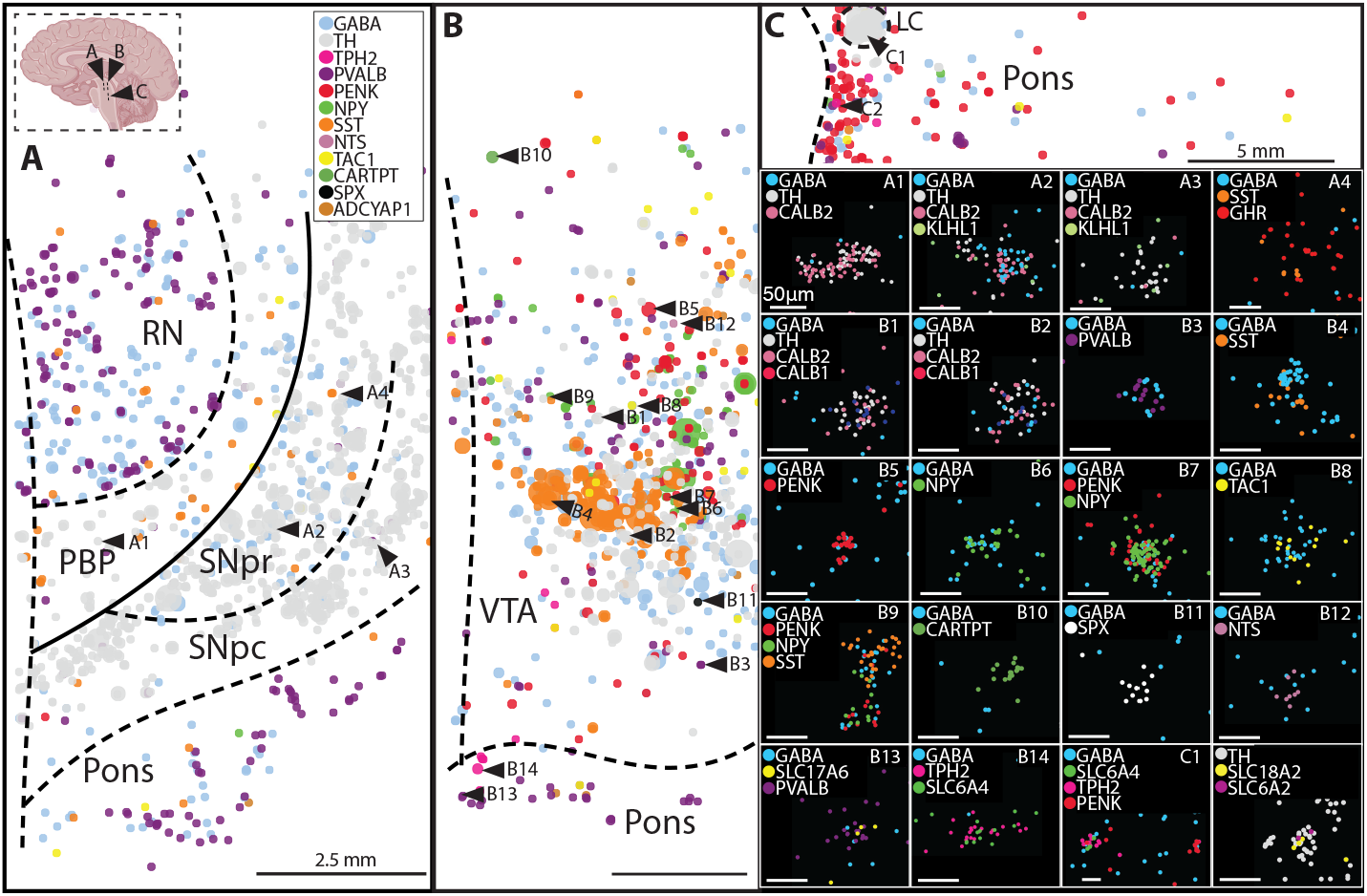
EEL-FISH reveals spatial distributions of gene expression in the midbrain and hindbrain. Panels A, B, and C each correspond to a tissue section. Cells are depicted by dots in each panel; the color and size of the dot correspond to a gene and that gene’s expression level, respectively. Dots overlap when multiple genes are expressed in the same cell; multiple genes expressed in a cell at similar levels will not be visible. Some cells are therefore indicated with an arrowhead and inset in A1 - C2, where each dot corresponds to a molecule rather than a cell. Tissue sections include (**A**) red nucleus (RN), parabrachial pigmented nuclei (PBP), substantia nigra pars reticulata (SNpr), substantia nigra pars compacta (SNpc), and pons; (**B**) ventral tegmental area (VTA) and pons; and (**C**) pons and locus coeruleus (LC).

The EEL data additionally mirrored our RNA-seq analysis of *splatter* neurons more broadly. Although we did not dissect VTA for sequencing, rodent VTA neurons resemble *splatter* neurons in their combinatorial expression of neurotransmitters, neuropeptides, and their receptors (*22*). Indeed, GABAergic and glutamatergic neurons in the pons and VTA combinatorially co-expressed a number of neuropeptides (Fig 4. B-C: B3-13,C1), and most *CALB1*+ dopaminergic cells in the VTA also expressed GABAergic genes (Fig. 4B: B1,B2). Some neuronal populations exhibited specific spatial distributions: SST+ GABAergic cells crossed the VTA axially, whereas *NPY*+ GABAergic and *PENK*+ GABAergic cells distributed sagittally (Fig. 4B, Fig. S6B). Serotonergic neurons scattered along the midline in the VTA and the pons (Fig. 4B: B14, Fig. 4C: C1). *TAC1*+ GABAergic neurons were instead distributed broadly throughout the tissue (Fig. 4B: B8). Notably, some populations were very rare. A spatially isolated cell situated in the substantia nigra *pars reticulata* co-expressed *GHR* and *SST* (Fig. 4A: A4). Fewer than three cells highly expressed peptides like *ADCYAP1, CARTPT, NPX*, or *NTS* (Fig 4B: B10-12; Fig. S6B). Our results therefore imply not only that *splatter* neurons encompass both broadly distributed and spatially restricted cell types, but also that their complexity reflects under-sampling. Cells across the midbrain and hindbrain are defined by combinatorial neuropeptide and neurotransmitter expression, and specific combinations might be observed in few cells that are difficult to sample with single-nucleus RNA sequencing from single or few donors.

We next analyzed the non-neuronal cells in our dataset, which included vasculature, immune cells, and non-neuronal neuroepithelial cells called glia that support brain function in diverse ways: astrocytes maintain brain homeostasis; oligodendrocytes myelinate neurons to facilitate electrical impulses. Three superclusters corresponded to the oligodendrocyte lineage: eight *oligodendrocyte* clusters, six *committed oligodendrocyte precursors* (*COPs*), and three *oligodendrocyte precursor cells* (*OPCs*). One *oligodendrocyte*, four *COP*, and one *OPC* cluster were likely low quality or doublets and excluded from further analysis (#41-43, 36, 75) The remaining clusters were projected onto a single UMAP embedding that mirrored lineage differentiation (Fig. 5A). The dendrogram grouped eight *oligodendrocyte* clusters into two major types that expressed either *OPALIN* or *RBFOX1*, as previously described (Type 1 and 2, respectively, Fig. 5B-C; Fig. S7A; Table S3) (*23*). *RBFOX1*+ oligodendrocytes are thought to represent a mature end state, consistent with their UMAP position and perhaps our observation that more genes are upregulated than downregulated in these cells (6,174 and 5,128 genes respectively). One distinct *oligodendrocyte* cluster largely derived from a single dissection: 88% of the cells in this cluster were from the hypothalamic mammillary nucleus in one donor. Moreover only two of the top 20 enriched genes were protein coding (*MDFIC2, COL10A1*), and neither gene was highly specific. This cluster therefore likely resulted from donor-specific or quality-related effects (Fig. S7B-F). The remaining oligodendrocytes were smoothly distributed along the Type 1-to-Type 2 manifold.

**Figure 5.**
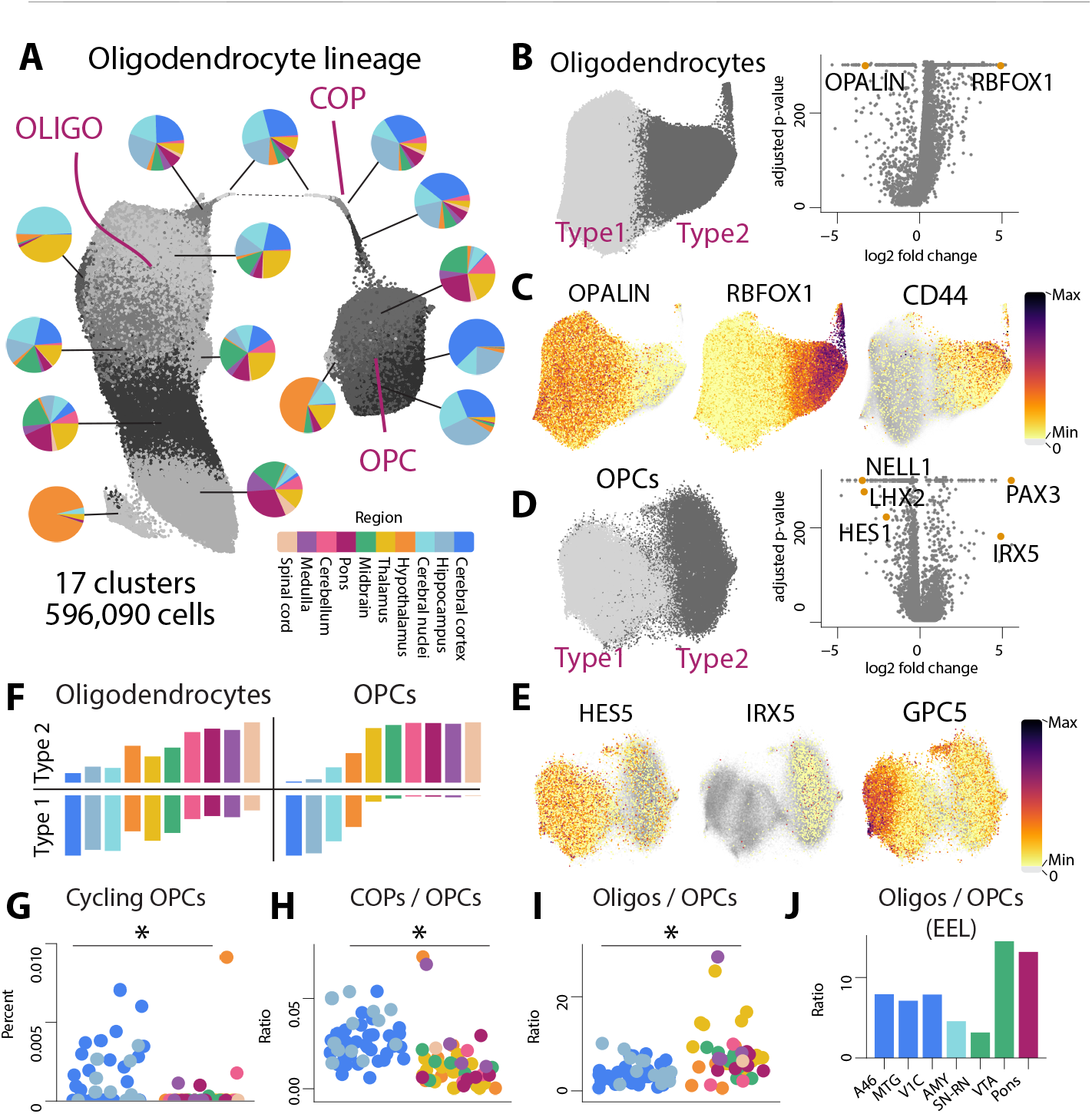
Oligodendrocyte-lineage cells differ between the telencephalon and the rest of the brain. (**A**) Oligodendrocytes, COPs, and OPCs are arranged on a UMAP embedding. Cells are grayscale-colored by cluster. Pie charts summarize each cluster’s regional composition according to the legend. (**B**) A dendrogram of oligodendrocyte clusters was cut to identify two transcriptomic types. Type 1 and Type 2 cells are colored on the t-SNE (left). Volcano plot shows genes differentially expressed between Type 1 and Type 2 with q-value less than 1e-5 and mean expression greater than 0.02 (right). (**C**) Cells are colored by gene expression. (**D**) Same as (B) but for OPCs. (**E**) Same as (C). (**F**) Bar heights represent the proportions of Type 1 and Type 2 (bottom and top, respectively) oligodendrocytes and OPCs (left and right, respectively) found in each region. Colors correspond to (A). (**G-I**) Each dot represents a dissection colored by its region of origin as in (A). Dots on the left are telencephalic dissections; dots on the right are from the rest of the brain. Dots within each group are jittered randomly along the x-axis for visualization. Dot height represents a dissection’s (G) percentage of cycling OPCs, (H) ratio of COPs to OPCs, or (I) ratio of oligodendrocytes to OPCs. Stars indicate a statistically significant difference between the two groups (Mann-Whitney, p-value < 1e-4). The y-axis in (G) was trimmed to 0.01 for visualization; one telencephalic dissection contained 0.04% cycling cells. (**J**) Bar heights represent the ratios of oligodendrocytes to OPCs in each tissue section profiled with EEL.

Because oligodendrocyte cells are relatively homogenous and therefore not necessarily well characterized by clustering analysis, we used Latent Dirichlet Allocation (LDA) to identify transcriptional modules that varied across these cells. LDA is most commonly used to identify groups of words—referred to as “topics”—that frequently appear together in text documents. The method analogously reveals co-expressed genes in single-cell data (*24*). We fit an LDA model for 15 topics (k=15) to the *oligodendrocyte* data and identified representative genes for each topic (Fig. S7G-L; Methods). Mitochondrial genes represented Topic 3, indicating low-quality cells that we did not detect with other metrics. Four other topics peaked successively along the Type 1-to-Type 2 manifold and thus resembled intermediate stages along a maturation axis; correspondingly, these topics were lead by a topic well-represented by *OPALIN* and ended with a topic well-represented by *RBFOX1* (Fig. S7I-L). LDA additionally enables the user to score new cells for a given gene topic. Because we did not observe *Rbfox1* expression in mouse oligodendrocytes, we scored myelin-forming and mature mouse oligodendrocytes for the *RBFOX1*-correlated Topic 6 (*1*). A subset of primarily non-telencephalic oligodendrocytes (MOL3 and some MOL2 cells) scored highest for Topic 6 (Fig. S7M). Among the genes most correlated with Topic 6 in mice, *VAT1L, NKX2-8* (mouse *Nkx2*-9), and *CD44* were strikingly specific in both species, suggesting these genes are conserved markers of Type 2 oligodendrocytes (Fig. 5B; Fig. S7N). Overexpression of the hyaluronan receptor *CD44* in oligodendrocytes promotes demyelination, which might signify myelin turnover (*25*).

The UMAP and nearest-neighbor graph suggested that the three *COP* clusters constituted a lineage trajectory linking OPCs to mature oligodendrocytes. In contrast, the four *OPC* clusters were characterized by striking regional specificity rather than maturation state (Fig. 5A; Fig. S7O). We used the cluster dendrogram to group OPCs into two major types and found each enriched within and outside the telencephalon (Type 1 and Type 2, respectively; Fig. 5D). These populations likely correspond to the *NELL1*+ cortical and *PAX3*+ spinal-cord populations described in a recent preprint (*26*). Type 1 and Type 2 OPCs differentially expressed over 155 genes with a greater than three-fold change in either population, including axon-guidance molecules, region-specific transcription factors, and Notch signals (Fig. S8A; Table S4). Although our previous work identified only one OPC cluster in the adolescent mouse, experimental differences likely explain the discrepancy; we sampled fewer cells in mice using a single-cell chemistry with lower sensitivity. We reanalyzed the data and found that type-specific markers *Hes5* and *Irx5* were expressed mutually exclusively in mouse OPCs (Fig. S8B). This observation highlights that OPCs exhibit relatively little transcriptomic variation—few principal components explained most of their variance (Fig. 3D)— but are not homogeneous across the brain.

We accordingly further assessed the four *OPC* clusters (Fig. S8C). One Type 2 cluster was 54% mammillary nucleus, reminiscent of an oligodendrocyte cluster (Fig. S8D-F). Although all three donors contributed to this cluster—207, 464, and 257 cells each—only *FMO3* of the top 20 enriched genes was protein-coding, consistent with a possible quality effect. Clustering within Type 1 cells appeared to be driven by two expression gradients that prevented our analysis pipeline from merging the clusters. Although the first gradient was correlated with total UMIs per cell, not all genes were uniformly upregulated in larger cells. Telencephalic-specific transcription factor *FOXG1* was evenly expressed across Type 1 OPCs, whereas the secreted protein *NELL1* and cell-adhesion molecule *CNTN5* increased smoothly (Fig. S8G-J). The second gradient was defined by the proteoglycan *GPC5*, previously identified as a marker for grey-matter astrocytes (Fig. 5E). OPC heterogeneity within the telencephalon might therefore reflect interactions with the local environment.

Because OPCs were distinct inside and outside the telencephalon, we examined more closely the regional distributions of oligodendrocytes. Type 1 oligodendrocytes predominated in the telencephalon, whereas the proportion of Type 2 oligodendrocytes progressively increased toward the brain’s more posterior regions (Fig. 5F). Because Type 1 oligodendrocytes were also putatively less mature than Type 2, we wondered whether the differential distributions of OPCs and oligodendrocytes along the anterio-posterior axis partly reflected different rates of oligodendrocyte turnover inside and outside the telencephalon. Telencephalic OPCs were indeed more proliferative than non-telencephalic OPCs (Fig. 5G). We therefore investigated the ratios of *OPC, COP* and *oligodendrocyte* populations across dissections, reasoning these ratios would reflect relative turnover rates. We found an elevated ratio of COPs to OPCs in the telencephalon, indicating more frequent initiation of differentiation (Fig. 5H). The ratio of oligodendrocytes to OPCs was instead higher in the brainstem, indicating lower turnover of mature cells relative to the progenitor population (Fig. 5I). EEL data confirmed the brain stem’s elevated ratio of oligodendrocytes to OPCs, except for a section that originated from an anterior position in the substantia nigra and red nucleus (SN-RN, Fig. 5J). Taken together, these observations suggest that telencephalic OPCs are more proliferative, differentiate more frequently, and form a larger population relative to oligodendrocytes. Elevated OPC proliferation might therefore more frequently replenish the oligodendrocyte population in the telencephalon, lowering the resident proportion of more mature Type 2 oligodendrocytes.

Whereas oligodendrocytes likely occupied a continuum of maturing states, mature astrocytes exhibited more sharply delineated heterogeneity across the brain. A GO-enrichment analysis of the most variable genes in these cells revealed synaptic-transmission, axon-guidance, and extracellular-matrix terms, highlighting that astrocytes are specialized to interact with nearby neurons and their extracellular environment (Fig. S9A). Two major astrocyte types are present in the mouse brain: a telencephalic and a non-telencephalic type, each of which additionally contains grey-matter (*Gfap*-negative) and white-matter (*Gfap*-positive) astrocytes. The cortex additionally contains interlaminar astrocytes that are specific to Layer 1 (*2*). Our clustering analysis showed that this molecular architecture is conserved in human astrocytes. We identified 13 *astrocyte* subpopulations that organized into a telencephalic and non-telencephalic type (Type 1 and 2, respectively), each containing *GFAP* low and high populations (Fig. 6A-B; Fig. S9B-D). Type 1 included the expected cortical populations: *WIF1*+ grey-matter, *TNC*+ white-matter, and *LMO2*+ interlaminar astrocytes. The latter appeared especially distinct on the embedding (Fig. 6C).

**Figure 6.**
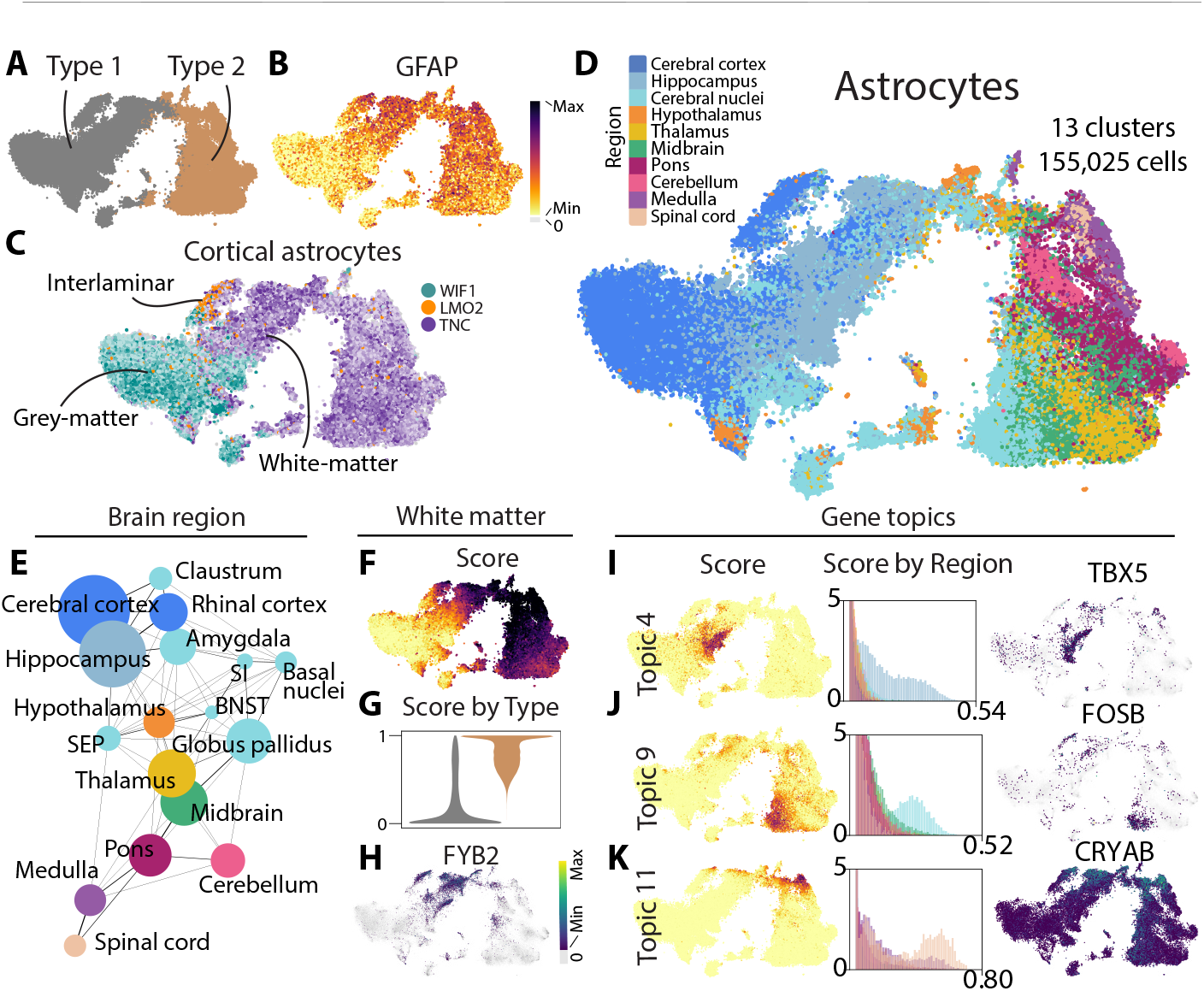
Astrocytes exhibit transcriptomic diversity across brain regions. (**A**) A dendrogram of astrocyte clusters was cut to identify two transcriptomic types colored on the t-SNE. (**B**) Cells are colored by gene expression. (**C**) Cells are colored by expression levels of the gene they express most highly among those in the legend. Darker color indicates higher expression; gray indicates no expression. Each gene is enriched in one of the cortical astrocyte types indicated on the t-SNE. (**D**) Cells are colored by their regions of origin. (**E**) A graph represents astrocyte similarities across the brain. Most nodes represent brain regions; cerebral nuclei are broken up into finer regions or dissections. Node size is proportional to the number of cells from each region or dissection. The weight of an edge between two nodes is proportional to the percentage of cells in the regions or dissections that are neighbors. Abbreviations include: BNST = bed nucleus of the stria terminalis; SEP = septal nuclei; SI = substantia innominata and nearby nuclei. (**F**) Cells are colored by their scores for a gene topic that mirrors *GFAP* expression and therefore likely reflects white-matter association. (**G**) Violin plots summarize score distributions for Type 1 and Type 2 astrocytes colored as in (A). (**H**) Cells are colored by gene expression. (**I-K**) Cells are colored by their scores for each gene topic (left). Histograms summarize the score distributions for each region, colored as in (C) (middle). Cells are colored by expression of a topic-representative gene (right).

We observed additional regional heterogeneity within both Type 1 and Type 2 astrocytes (Fig. 6A). Astrocyte clusters were distributed uniquely across most brain regions, with astrocyte composition most similar across neighboring developmental and anatomical compartments (Fig. S9E-F). The cortex, for example, contained two gray-matter clusters (#52 and #54), one enriched in neocortex and another in allocortex, mirroring differences across cortical neurons. Whereas the former was mostly specific to the cortex, the latter was also found in the hippocampus and amygdala. The two other cortical clusters—interlaminar (#58) and fibrous (#61) astrocytes—were more evenly distributed across the entire cortex, perhaps reflecting less specialization for specific neuronal circuitry. Hippocampal astrocytes were found not only in all four cortical clusters, but also two largely hippocampus-specific clusters (#53 and #57). Astrocytes in the cerebral nuclei showed especially high diversity, reflecting the diversity of neuronal circuitry this brain region encompasses (Fig. S9G). Whereas most amygdala astrocytes clustered with allocortical astrocytes (#54), striatal astrocytes formed their own PENK+ cluster highly distinct on the t-SNE (#55). Substantia innominata cells were most similar to striatum, but the claustrum to cortex and the septal nuclei to the hypothalamus. BNST astrocytes clustered with amygdala, hippocampal, and striatal cells. Most surprisingly, globus pallidus astrocytes resembled midbrain and thalamus astrocytes and therefore represented the only major telencephalic Type 2 cells. Other Type 2 cells derived from the diencephalon, midbrain, and hindbrain. Hindbrain astrocytes were largely distributed across two clusters, one dominated by medulla and spinal cord, and the other containing cerebellar astrocytes. Pontine astrocytes were found in both of these clusters. Importantly, our clustering analysis did not fully discriminate the regional heterogeneity we observed on the t-SNE. Some midbrain, hypothalamus, and thalamus cells that co-clustered with pontine astrocytes are therefore likely transcriptionally distinct (compare Fig. 6D and Fig. S9B). Transcriptomic variation among grey- and white-matter astrocytes possibly confounded the analysis, particularly because astrocyte clusters were distinguished mostly by combinatorial gene-expression gradients rather than binary markers (Fig. S9H).

To distinguish heterogeneity that our clustering analysis failed to capture, we again used LDA to identify gene topics differentially distributed across astrocytes. First, we fit an LDA model with only two topics (k=2) and recovered a topic that mirrored *GFAP* expression(Fig. 6B,F). We used this topic to score astrocytes for their likelihood to be white-matter associated. The “white-matter score” fell in a gradient across both Type 1 and Type 2 astrocytes, as well as within individual clusters like interlaminar astrocytes (#5). Notably, the score was differentially distributed across Type 1 and Type 2 astrocytes (Fig. 6F,G), which might suggest that we sampled mostly white-matter astrocytes outside the telencephalon. Consistent with this hypothesis, white-matter scores decreased along the antero-posterior axis in a trend that mirrored expected myelination levels (Fig. S10A). However, these types also appeared transcriptionally diverse across regions; FYB2, for example, was highly specific to Type 1 white-matter astrocytes (Fig. 6F).

We next fit an LDA model with 30 topics (k=30). Although many topics correlated with individual clusters, we also identified gene topics that scored highly within and across clusters. Topics 5 and 29 appeared to differentiate Type 2 grey- and white-matter astrocytes (Fig. S10B). Topic 22 instead revealed an expression gradient of growth factor *HGF* within cortical grey-matter astrocytes (Fig. S10D), and Topic 4 was best represented by the transcription factor *TBX5*, which is highly enriched in hippocampus-specific astrocytes (Fig. 6I, clusters #53 and #57). Notably, Topic 9 was elevated in the globus pallidus and well represented by high *FOS* and *JUN* expression (Fig. 6J). *FOS* is activated downstream of a range of cellular processes including ischemia, which would agree with our observation that globus pallidus and substantia nigra dissections contained unusually high numbers of microglia (Fig. S1B). Pathological conditions might therefore have driven our classification of many globus pallidus astrocytes as Type 2. We identified two other topics best represented by genes previously characterized as potential markers for reactive astrocytes (*27*). Topic 9 was elevated in all fibrous astrocytes and correlated with *NR4A1* and *CRYAB* expression (Fig. 6K). Topic 8 was most elevated in a few tiny groups on the embedding and correlated with inflammatory-response genes *CHI3L1, SERPINA3*, and *CCL2* (Fig. S10E). Together, these results emphasize both the regional heterogeneity of astrocytes and the diverse molecular programs with which they react to their environments.

Here we have presented the first survey of transcriptomic cell-type diversity across the entire human brain. Our study highlights some of the challenges associated with profiling human tissue: we likely observed pathological states like the response to ischemia (e.g. astrocytes, Fig. 6), and clusters were not always reproducible across donors, partly due to the difficulties of replicating anatomical dissections (e.g. hypothalamus, Fig. 2). How cell types vary across donors will be an exciting avenue for future research. Our study also highlights a remarkable gap in our current understanding of neuronal diversity. The telencephalon was largely populated by abundant, relatively homogeneous cell types like layer-specific excitatory neurons, cortical interneurons, and MSNs. In contrast, the molecular architecture of *splatter* neurons—encompassing many tiny, well-defined clusters and larger highly heterogenous clusters—likely indicates that we are highly under-sampled in regions like the hypothalamus, midbrain, and hindbrain, which control innate behaviors and autonomous physiological functions. Our previous work did not uncover equivalent complexity in the mouse, but here we have identified subclusters that are likely conserved across species, suggesting that deeper sampling will reveal similar neuronal complexity in the mouse. Sampling depth however might not fully explain the striking complexity of these neurons; *splatter* neurons uniquely comprised both GABAergic and glutamatergic neurons that did not separate on the embedding, and they expressed neuropeptides and other neuronal genes in unusually complex combinatorial patterns. Our results additionally underscore how strongly developmental origins shape adult transcriptomic types. Both astrocytes and OPCs exhibited telencephalic and non-telencephalic types that transitioned in the hypothalamus (compare Fig. S7O with Fig. S9E). Some superclusters reflected migratory cell types that appeared in multiple dissections, including some not previously observed. A thorough understanding of region-specific cellular diversity and its development will therefore be key for treating disease. For example, the unique composition of the telencephalon’s oligodendrocyte lineage might be relevant for diseases like multiple sclerosis. Our work therefore provides a critical foundation for exploring the diverse neural circuitry across the brain and its implications for human health.

## Acknowledgements

We are very grateful to members of the Linnarsson group for feedback and discussion. We especially thank Anna Johnsson for lab management, Egle Kvedaraite for immune-cell annotation, Elin Vinsland for fibroblast and vasculature annotation, and Ivana Kapustová for the schematic drawing of the human brain. We also acknowledge support from the National Genomics Infrastructure in Stockholm funded by Science for Life Laboratory, the Knut and Alice Wallenberg Foundation and the Swedish Research Council, and SNIC/Uppsala Multidisciplinary Center for Advanced Computational Science for assistance with massively parallel sequencing and access to the UPPMAX computational infrastructure. We thank the tissue procurement, tissue processing and facilities teams at the Allen Institute for Brain Science for assistance with the transport and processing of human brain specimens and the technology team at the Allen Institute for assistance with data management. The content of this article is solely the responsibility of the authors and does not necessarily represent the official views of the National Institutes of Health.

## Funding

European Molecular Biology Organization fellowship

ALTF 1015-2018 to KS

U.S. National Institutes of Health (NIH) grant U01 MH114812-02 to SL, ESL

Chan Zuckerberg Initiative and the Silicon Valley (NDCN 2018-191929) to EA, SL

Knut and Alice Wallenberg Foundation (2015.0041, 2018.0172, 2018.0220 to SL; 2018.0232 to EA)

Swedish Foundation for Strategic Research (SB16-0065) to SL, EA

Torsten Söderberg Foundation to SL

European Union grants: ERC-ADG 884608, H2020 874758 to EA

Vetenskapsrådet 2020-01426, Karolinska Institutet (StratRegen SFO 2018) and Hjärnfonden FO2019-0068to EA.

## Author contributions

Study design: SL, EL, RDH, TB, SD, KS

Tissue dissections and nuclei isolation: RDH, SD, MC, TC, ND, JG, CDK, JN, HT, AMY

RNA sequencing: LH, KS RNA data preprocessing: PL RNA data analysis: KS, SL

RNA data annotation: KS, AMA EEL experiments and analysis: AMA Dopaminergic analysis: KWL, EA RNAScope: RDH

Writing: KS, AMA, EA, KWL, RDH

Editing: KS, SL

## Competing interests

The authors declare that they have no competing interests.

## Data and materials availability

The data were generated for the BRAIN Initiative Cell Census Network (BICCN,

RRID:SCR_015820) and are available for download from the Neuroscience Multi-omics Archive (NeMO, RRID:SCR_016152) at: http://data.nemoarchive.org/biccn/grant/u01_lein/linnarsson/transcriptome/sncell/10x_v3/human/raw/. The annotated expression matrix, as well as code used for analysis and figure generation, is available on Github at linnarsson-lab/ adult-human-brain. Auto-annotations are available at linnarsson-lab/auto-annotation-ah.

## Methods

### Human postmortem tissue specimens

De-identified postmortem adult human brain tissue was obtained after receiving permission from the deceased’s next-of-kin. Tissue collection was performed in accordance with the provisions of the United States Uniform Anatomical Gift Act of 2006 described in the California Health and Safety Code section 7150 (effective 1/1/2008) and other applicable state and federal laws and regulations. The Western Institutional Review Board reviewed tissue collection procedures and determined that they did not constitute human subjects research requiring institutional review board (IRB) review. The collection and processing of human tissue was approved under Swedish law by the Swedish Ethical Review Authority (2019-03054).

Male and female donors 18–68 years of age with no known history of neuropsychiatric or neurological conditions were considered for inclusion in the study. Routine serological screening for infectious disease (HIV, Hepatitis B, and Hepatitis C) was conducted using donor blood samples and donors testing positive for infectious disease were excluded from the study. Specimens were screened for RNA quality and samples with average RNA integrity (RIN) values ≥7.0 were considered for inclusion in the study. Postmortem brain specimens were processed as previously described (dx.doi.org/10.17504/protocols.io.bf4ajqse). Briefly, coronal brain slabs were cut at 1 cm intervals, frozen in dry-ice cooled isopentane, and transferred to vacuum-sealed bags for storage at -80°C until the time of further use. To isolate regions of interest, tissue slabs were briefly transferred to -20°C, and the region of interest was removed and subdivided into smaller blocks on a custom temperature controlled cold table. Tissue blocks were stored at -80°C in vacuum-sealed bags until later use.

### Single-nucleus RNA sequencing

Nucleus isolation for 10x Chromium RNA sequencing was conducted as described (dx.doi.org/10.17504/protocols. io.y6rfzd6). Gating on DAPI and NeuN fluorescence intensity was described previously (2). NeuN+ and NeuN-nuclei were sorted into separate tubes and were pooled at a defined ratio after sorting. Sorted samples were centrifuged, frozen in a solution of 1X PBS, 1% BSA, 10% DMSO, and 0.5% RNAsin Plus RNase inhibitor (Promega, N2611), and stored at -80°C until the time of shipment on dry ice from the Allen Institute to the Karolinska Institute for 10x chip loading.

Immediately before loading on the 10x Chromium instrument, frozen nuclei were thawed in a 37°C water bath, spun down briefly, and pipetted several times to mix (dx.doi.org/10.17504/protocols.io.eq2lyrdpgx9k/v2). Nuclei were then quantified and loaded according to the 10X Genomics protocol, targeting 5000 cells. We aimed to sequence two replicates per sample. Nearly all samples were processed with the 10x Genomics V3 kit; 12 samples were sequenced with v3.1 (Table S1). Samples were first sequenced to a shallow depth (approximately 1000 reads/cell) on the Illumina NextSeq platform to validate sample concentrations. Samples were then sequenced to approximately 100,000 reads/cell on the Illumina NovaSeq platform. Sequencing saturation was examined for each sample using the preseq package (https://github.com/smithlabcode/preseq). Any samples that were not saturated to 60% were sequenced more deeply using preseq predictions.

### Data preprocessing

Sequencing runs were demultiplexed with cellranger mkfastq version 4.0.0 (10x Genomics) and filtered through the index-hopping-filter tool version 1.1.0 (10x Genomics). Unique molecular identifier (UMI) counts were determined using STARSolo version 2.7.10a with the following parameters:

- -soloFeatures Gene Velocyto
- -soloBarcodeReadLength 0
- -soloType CB_UMI_Simple
- -soloCellFilter EmptyDrops_CR %s 0.99 10 45000 90000 500 0.01 20000 0.01 10000
- -soloCBmatchWLtype 1MM_multi_Nbase_pseudocounts
- -soloUMIfiltering MultiGeneUMI_CR
- -soloUMIdedup 1MM_CR
- -clipAdapterType CellRanger4
- -outFilterScoreMin 30

Barcode whitelists were downloaded from the 10x Genomics website. Exonic, intronic, and ambiguous counts were summed for clustering analysis.

The reference genome and transcript annotations were based on the human GRCh38.p13 gencode V35 primary sequence assembly. However, we filtered the reference. Because our pipeline only counted reads that uniquely aligned to one gene, reads that aligned to more than one gene were lost. Related genes therefore exhibited few or zero counts. To minimize this loss, we discarded overlapping genes or transcripts that overlapped or mapped to other genes or non-coding RNAs’ 3’ UTRs, leaving only one of these transcripts in the genomic reference. We used BLAST to align the last 400 nucleotides (3’ UTR) of all protein-coding and non-coding transcripts to all other genes (maximum 4 mismatches, minimum alignment length 300 nucleotides). We resolved all the matches by the following procedure:

- Fusion genes were filtered based on their names: genes with names that contained both fusion genes [“gene1-gene2”] were discarded.
- Non-coding transcripts that matched a coding transcript were discarded.
- For overlapping transcripts where both transcripts were either coding or non-coding:

1. If one of the gene names matched the pattern “XX######.#” its transcript was discarded.
2. If one of the transcripts belonged to a gene with one or more transcripts that were already discarded during the procedure, it was discarded as well.
3. We discarded transcripts of the gene with fewest splice variants.
4. Otherwise, the transcript with a 3’ UTR that overlapped the other transcript was kept in the genomic reference
5. For paralogs, we mapped all related genes (that aligned to one another) and selected the paralog with most splice variants. All other highly similar paralogs were discarded. In special cases, we manually chose the gene.

Altogether we filtered 387 fusion genes, 1140 overlapping transcripts, 414 non-coding transcripts, 1127 coding paralogs, and 350 non-coding paralogs.

### Quality control

10x Chromium samples that captured more than 15,000 cells were not analyzed further (Table S1). All remaining samples were pooled with their replicates and analyzed with “cytograph qc” (commit 5f07e59768844e3c84fceb8fd7d1993059e13a7e), which uses a modified version of DoubletFinder to calculate a doublet score for each cell (28). Cells with fewer than 800 UMIs, fewer than 40% unspliced molecules, or a doublet score below 0.3 were removed from further analysis. These thresholds were determined by examining distributions across the dataset (Figure S1).

### Clustering

Cells were clustered using an updated version of the Cytograph package (github.com/linnarsson-lab/adult-human-brain,commit 22a83ebcf79a9fdcdff34beb0300aa1695d7039b) (1). The command “cytograph process” was run with configuration min_umis: 0, doublets_action: None, features: variance, remove_low_quality: False, remove_doublets: False, batch_keys: [“Donor”], clusterer: sknetwork. Default values were used for other parameters. In brief, principal component analysis (PCA) was performed on the most variable genes and used to construct a k-nearest neighbors graph, which provided input for Louvain clustering (scikit-network). Harmony with default parameters was used to integrate principal components across donors before constructing the graph (6).

This clustering pipeline was used at four levels of analysis: (1) All cells that passed QC were pooled into a single dataset and analyzed. This top-level clustering was split by the dendrogram into two groups that corresponded to either neuronal or non-neuronal cells, with one exception: lymphocytes clustered with neurons. (2) Each of these two groups was separately analyzed. (3) Paris clustering was used on each graph to find “superclusters” within each group for further analysis (Figure 1) (7). The Paris dendrogram was arbitrarily cut so that the superclusters mirrored large islands on the t-SNE. Each supercluster was separately analyzed to produce “clusters.” (4) Each of the 461 clusters was analyzed separately to produce “subclusters.”

At Level 3, additional cluster-level quality control was performed. A cluster was removed from further analysis if: (1) its mean percentage of mitochondrial UMIs was greater than the 99^th^ percentile mean (0.056); or (2) if the Cytograph pipeline tagged the cluster with two or more of the following auto-annotations: NEUR, ASTRO, OLIGO, OPC, MGL, ENDO, BERG, or VLMC. Auto-annotations and further information about the auto-annotation procedure are available at github.com/linnarsson-lab/ auto-annotation-ah. After removing these clusters, the remaining cells in each supercluster were re-clustered. This procedure was repeated until no clusters met these criteria.

Finally, clusters and subclusters at Levels 3 and 4, respectively, underwent a merging procedure (“cytograph merge”). A cluster that did not express at least one statistically enriched gene was merged with the nearest cluster on the t-SNE. In short, FDR-corrected p-values were calculated by comparing Cytograph’s enrichment scores with a null distribution obtained by permuting cluster labels across all cells in the supercluster.

A t-distributed stochastic neighbor embedding (t-SNE) and dendrogram were calculated for the final 461 clusters (Figure 1). The dendrogram was calculated based on Euclidean distances between cluster means for the 2,000 most variable genes. Cluster means were normalized to median cluster means across the dataset and then standardized (scikit-learn StandardScaler).

### Annotation

Superclusters were manually annotated based on the literature and on their regional composition. Clusters were not named with a single annotation but instead tagged with auto-annotations (github.com/linnarsson-lab/auto-annotation-ah) from four categories: neurotransmitters (auto-annotation-ha/Human_adult/Neurotransmission), neuropeptides (auto-annotation-ha/Neuropeptides), class (auto-annotation-ha/Human_adult/Class), and subtype (auto-annotation-ha/Human_adult/Subtype). Neuropeptide-related tags were manually compiled based primarily on the NeuroPep database (29). Subtype auto-annotations included manually selected cell types and states previously described in the literature. To identify cortical cell types, we also transferred labels from a single-cell analysis of the middle temporal gyrus (MTG) (30). We performed PCA (50 components) on our full dataset, trained a random forest classifier (scikit-learn, class_weight=‘balanced’, max_depth=50) on the MTG labels, and then predicted labels for all cells. We labeled each cluster with the mode of its constituent cells if two conditions were met: more than 0.8 of predicted labels matched the mode, and the mean probability of these predictions was greater than 0.8. Clusters that did not meet these criteria were labeled N/A.

### Gene ontology-enrichment analysis

The *gget* package (https://github.com/pachterlab/gget) was used to perform all gene-ontology enrichment analysis (function “enrichr” with database=“ontology”) (31).

### Assessing supercluster complexity with PCA

We used a permutation test to determine the number of principal components necessary to describe each supercluster, reasoning that this value would be proportional to the supercluster’s complexity. We therefore performed PCA twice on each supercluster (scikit-learn PCA, n_comps=50): once on the original expression matrix, and once on a permuted expression matrix, where the expression values of each gene were randomly redistributed across cells. We compared the percentage of variance that each principal component explained (attribute “explained_variance_ratio_”) in the original and unpermuted matrix. We considered the first component where explained_variance_ratio_ was similar between the two analyses (less than 10% higher in the unpermuted analysis than in the permuted) to represent the optimal number of components to describe the supercluster (Figure 3D, left). For each supercluster, we calculated an additional value using only the unpermuted analysis: the ratio between the percentages of variance explained by the first and second components (Figure 3D, right). We reasoned that this value illustrates how quickly the explained_variance_ratio decays across principal components. The script is available as permuted_pca.py at github.com/linnarsson-lab/adult-human-brain.

### RNAscope Multiplex fluorescent in situ hybridization (mFISH)

Fresh-frozen human postmortem brain tissues were sectioned at 10 μm onto Superfrost Plus glass slides (Fisher Scientific). Sections were dried for 10 minutes at -20°C and then vacuum sealed in plastic slide boxes and stored at -80°C until use. The RNAscope multiplex fluorescent v2 kit was used per the manufacturer’s instructions for fresh-frozen tissue sections (ACD Bio), except that fixation was performed for 60 minutes in 4% paraformaldehyde in 1X PBS at 4°C and protease treatment was shortened to 15 minutes. Sections adjacent to those used for RNAscope were fixed for 15 minutes in 4% paraformaldehyde in 1X PBS, briefly washed in PBS, and stained with NeuroTrace 500/525 Green Fluorescent Nissl Stain and DAPI to aid in region localization. Nissl-stained sections were imaged using a 10X objective on a Nikon TiE fluorescence microscope equipped with NIS-Elements Advanced Research imaging software (v4.20, RRID:SCR_014329). RNAscope sections were imaged using a 40X oil immersion lens on the same microscope. For mFISH experiments, positive cells were called by manually counting RNA spots for each gene. Cells were called positive for a gene if they contained ≥ 5 RNA spots for that gene. Lipofuscin autofluorescence was distinguished from RNA spot signal based on the larger size of lipofuscin granules and broad fluorescence spectrum of lipofuscin. Staining for the probe combination was repeated with similar results on at least 3 separate sections from one human donor. Images were assessed with FIJI distribution of ImageJ v1.52p and Nikon NIS-Elements imaging software. Probes used were Hs-OTX2-C3 (#484581-C3), Hs-SOX14-C2 (#1055351-C2), and Hs-CASR (#411931) from ACD Bio.

### Extended analysis of dopaminergic neurons

To identify additional dopaminergic subtypes, subclusters 1870, 1871, 1873, and 1876 were reanalyzed separately as described under “Clustering.”

To compare our data with a recently published census of midbrain dopaminergic neurons from Kamath et. al (4), we trained a random forest classifier on nuclei from their dataset annotated as dopaminergic neurons from control samples with non-neurological disorders (Donor IDs: 3298, 3322, 3345, 3346, 3482, 4956, 5610, 6173). We trained the classifier with features that Cytograph selects when run with the option to select cluster-enriched rather than variable genes (default parameters except n_factors: 20 and features: enrichment). The scikit-learn RandomForestClassifier was regularized with the parameters max_features and min_samples_leaf. Five-fold cross-validation was performed on data from Kamath et al. with scikit-learn StratifiedShuffledSplit for max_features 0.3, 0.4, 0.5 and min_samples_leaf between 3 to 50. We selected max_features=0.3 and min_samples_leaf=4 to transfer labels to our data; oob_score was set true, and other parameters were left as default.

### EEL-FISH

EEL-FISH was performed as recently described (15). Donor H18.30.002 was used for the V1C sample; H20.30.002 for the AMY, MTG and A46 samples; H19.30.001 for the pons sample; and H18.30.001 for the SN-RN and VTA samples. For all figure panels, genes were only plotted for an individual cell if the cell contained more than two molecules of that gene. Expression below this threshold was considered experimental noise.

### Differential-expression testing

Differential-expression testing was performed with the diffxpy package, version 0.7.4+21.g12f1286. Datasets were subset before analysis; 25,000 cells were randomly selected from each test group. Cells were normalized to the median total UMI count across the dataset and then used as input for a Mann-Whitney rank test (function “pairwise” with test=“rank”).

### Topic modeling with Latent Dirichlet Allocation

We used tomotopy version 0.12.2 to perform topic modeling with Latent Dirichlet Allocation (LDA). In brief we trained a model with a specified number of topics “k” until perplexity stabilized (function “LDAModel” with parameters alpha=50/k, eta=0.1). We trained with no more than 200,000 cells and all protein-coding genes in the genome that were expressed in at least 100 cells and fewer than 60% of all cells. Some topics were donor-specific or quality-specific, e.g, mitochondrial or ribosomal, and some did not have an apparent biological meeting. We therefore identified topics of interest manually. We also identified representative genes for each topic by using the topic probabilities reported for each gene (function “get_topic_word_dist”). We filtered genes by specificity (topic probabilities for each gene normalized by that gene’s probability across all topics) and then sorted the remaining genes by their unnormalized probabilities. A script is available as optimize_lda.py at github.com/ linnarsson-lab/adult-human-brain.

**Supplementary Figure 1.**
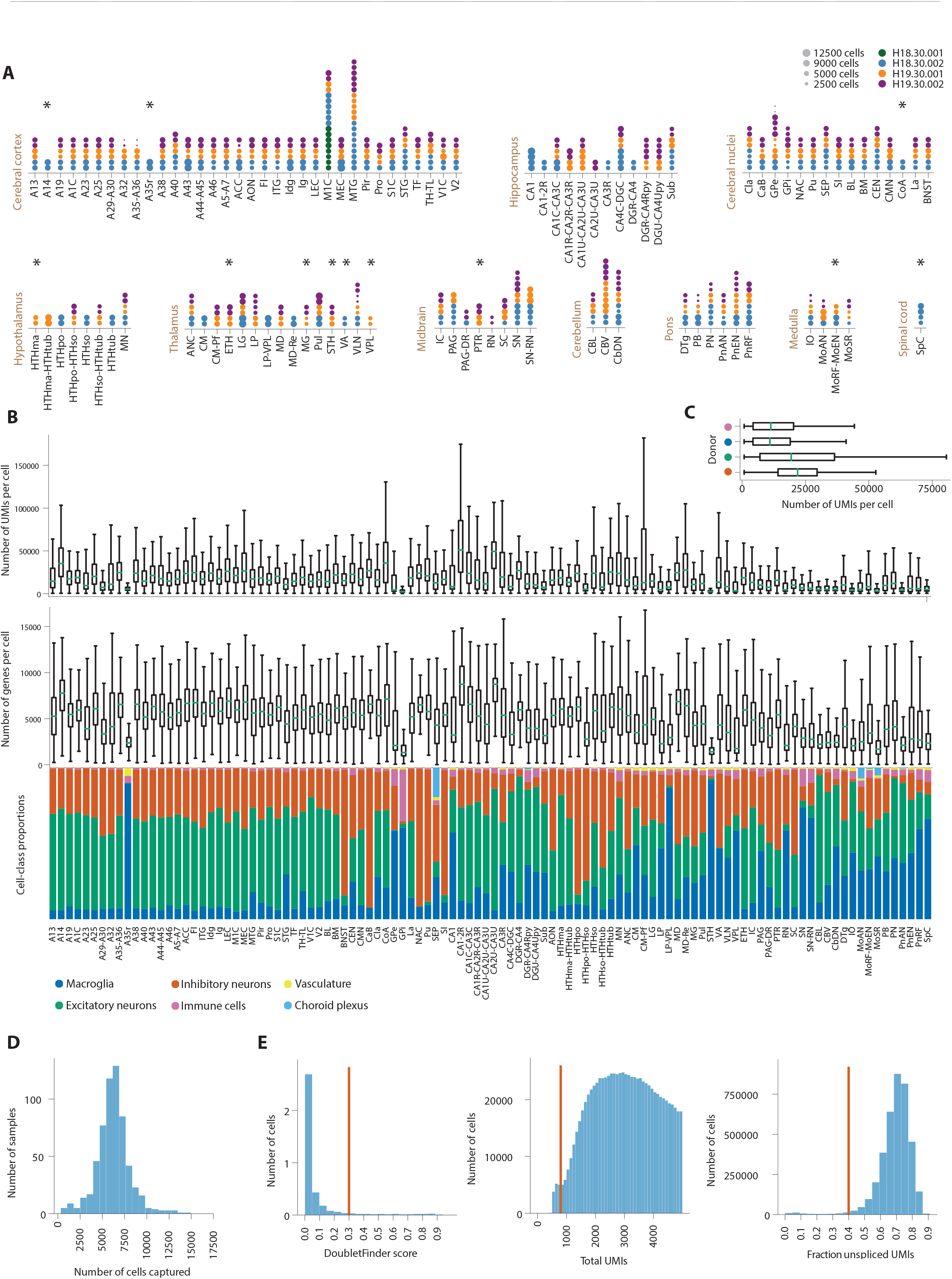
Overview and quality of the dataset. (**A**) Anatomical locations dissected within each brain region are shown along the x-axes. These locations are hereafter referred to as “dissections.” For each dissection, vertical dots represent 10X Genomics samples that passed quality control. Dot size corresponds to the number of cells obtained from each sample, as indicated by the legend. Dot color represents the sample’s donor of origin. Samples that were not replicated across all three donors are starred. Some samples that appear not to have been replicated might have been pooled differently across donors, such as red nucleus (RN), which was pooled with substantia nigra (SN) in two donors but not the third. (**B**) For each dissection, box plots without fliers summarize the distributions of total unique molecular integer (UMIs, top) and total genes (middle) per cell. Stacked bar plots summarize the proportions of cell classes sampled from each dissection (bottom). Each bar is colored by class as indicated by the legend. (**C**) For each donor, box plots without fliers summarize the distributions of total UMI counts per cell. (**D**) A histogram summarizes the distribution of cells captured per 10X Genomics sample. (**E**) Histograms summarize the distributions of metrics used to perform quality control on the complete dataset: DoubletFinder score per cell, total UMI count per cell, and percentage of unspliced molecules in each cell. Orange bars in each plot indicate the threshold used to remove low-quality cells.

**Supplementary Figure 2.**
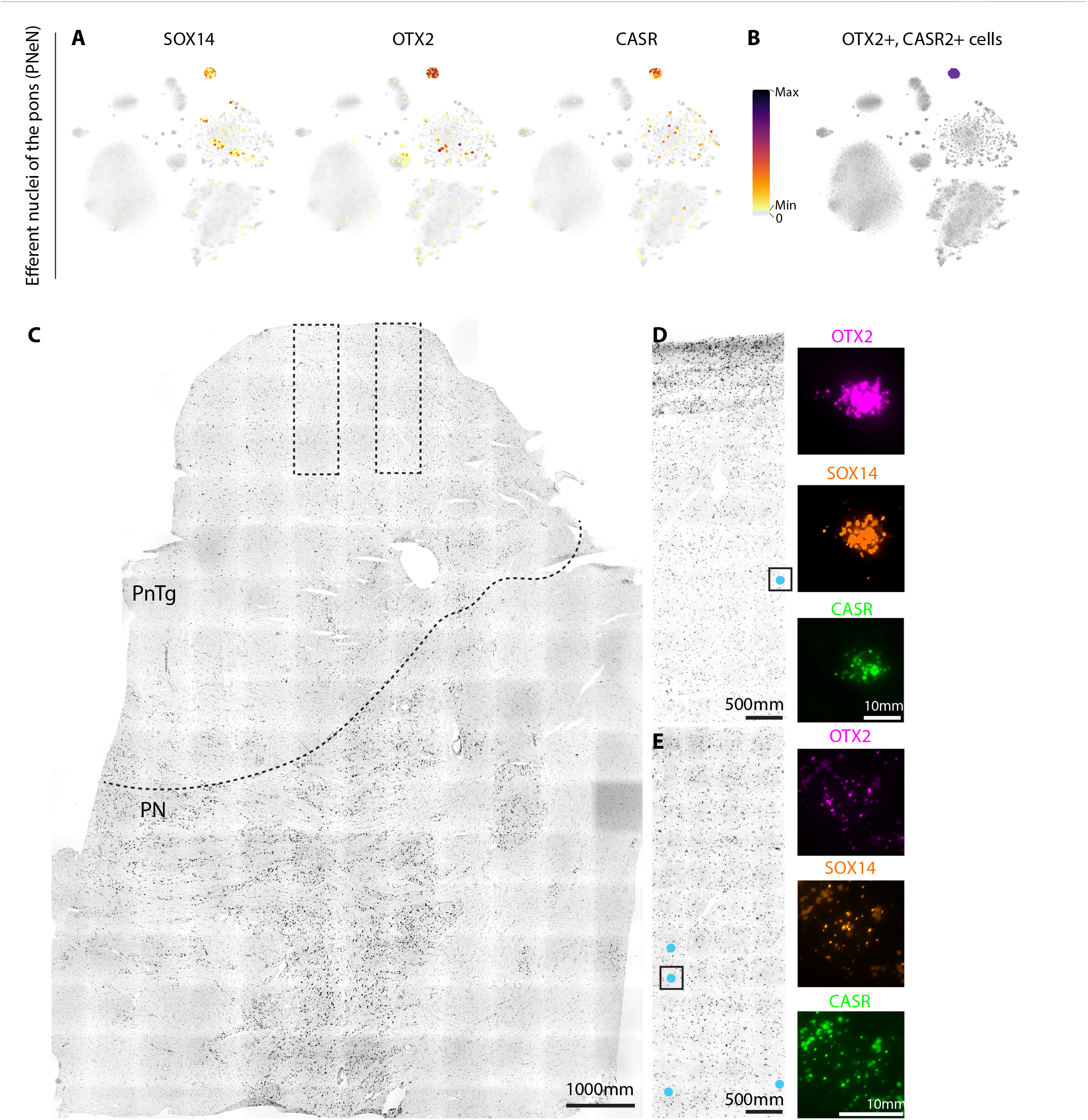
Midbrain-derived inhibitory neurons localize in the pons. **(A)** Cells from the efferent nuclei of the pons (PnEN) are arranged by t-SNE and colored by their expression of marker genes for midbrain-derived inhibitory neurons. **(B)** Cells on the t-SNE are colored purple if they express both OTX2 and CASR. **(C)** A Nissl-stained overview of a section is adjacent to one processed with RNAscope. Pontine nucleus (PN) and the pontine tegmentum (PnTg) that contains PnEN are delineated. Two boxes approximate the regions imaged in D and E. (**D-E**) On the left-hand side, the cyan circles mark cells that are positive for midbrain-inhibitory markers OTX2, SOX14, and CASR. Black boxes indicate the cells shown in the fluorescent images on the right-hand side.

**Supplementary Figure 3.**
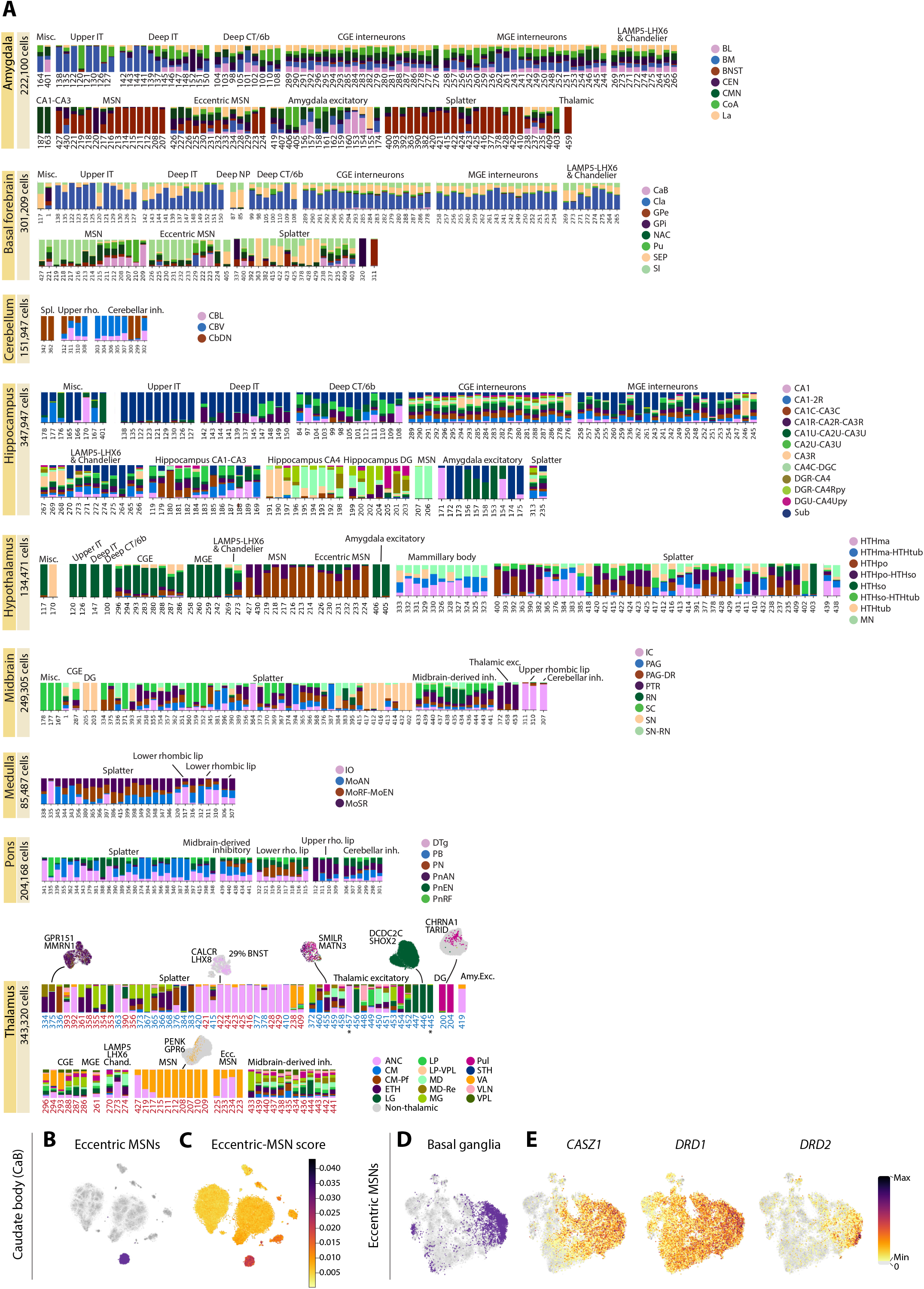
Superclusters and clusters are differentially distributed across brain regions. **(A)** Bar plots represent neuronal cluster distributions for each region as in Fig. 2B. **(B)** Cells from the caudate body (CaB) in the striatum are arranged by t-SNE; e*ccentric MSN* cells are purple. **(C)** CaB cells are colored by a score that represents the proportion of total UMIs that derive from the top 100 markers listed for mouse eccentric MSNs by Saunders et al. **(D)** *Eccentric MSNs* on the t-SNE are purple if they derive from the basal ganglia. **(E)** Cells are colored by gene expression.

**Supplementary Figure 4.**
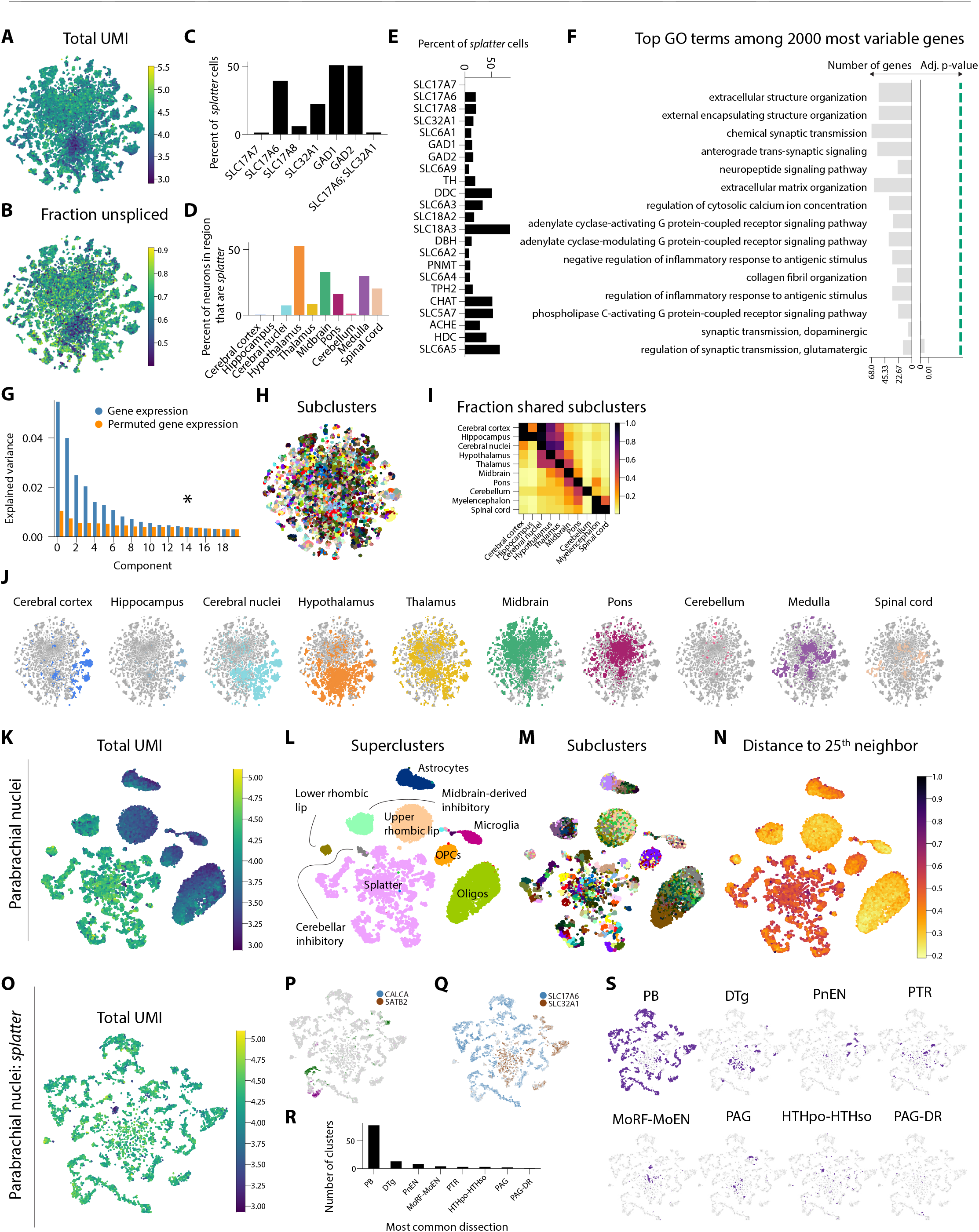
Splatter neurons are highly complex and undersampled. **(A)** Splatter cells are colored on the t-SNE by log10 of their total UMI counts. (**B**) Cells on the t-SNE are colored by fraction of unspliced UMIs. (**C**) The height of each bar indicates the percentage of *splatter* neurons that express the gene (or gene combination) shown on the x-axis. (**D**) The height of each bar indicates the percentage of neurons from each brain region that are *splatter* neurons. (**E**) The length of each bar indicates the percentage of *splatter* neurons that express the neurotransmitter shown on the y-axis. (**F**) A bar plot shows the most enriched gene-ontology terms among the 2000 genes most variable across *splatter* neurons. For each term, bar lengths represent the number of variable genes that overlapped with the term (left) and p-value of the enrichment (right). Green line indicates 0.05. (**G**) An example plot illustrates how the values shown in Fig. 3D were calculated for one supercluster. Bar height represents the fraction of total variance explained by each principal component along the x-axis. Principal components were calculated using both the original gene-expression matrix (blue) and a matrix in which each gene’s expression was permuted across cells (orange). The star indicates the value that is plotted left in Fig. 3D: the component where the fraction of explained variance in the original dataset is less than 10% greater than that of the permuted dataset. (**H**) *Splatter* cells on the t-SNE are colored by subcluster. (**I**) Each row in the matrix represents the fractions of subclusters from each region that are shared with regions along the x-axis. To be considered shared, subclusters contained at least five cells from both regions. (**J**) *Splatter* cells on each t-SNE are colored if they derive from the indicated region. (**K**) Cells from the parabrachial nuclei are colored on the t-SNE by log10 of their total UMI counts. (**L**) Cells are colored by supercluster, labeled on the panel. (**M**) Cells are colored by subcluster. (**N**) Cells are colored by Euclidean distance to their 25^th^ nearest neighbors. Distances were normalized by the maximum distance in the dataset. (**O**) *Splatter* neurons from the parabrachial nuclei are colored on the t-SNE by log10 of their total UMI counts. (**P-Q**) Cells are colored by expression levels of the gene they express most highly among those in the legend. Darker color indicates higher expression; gray indicates no expression. (**R**) Bar height indicates the number of parabrachial *splatter* subclusters for which the dissection along the x-axis was the most common dissection. (**S**) Cells are colored purple if they derive from a subcluster predominated by the indicated region.

**Supplementary Figure 5.**
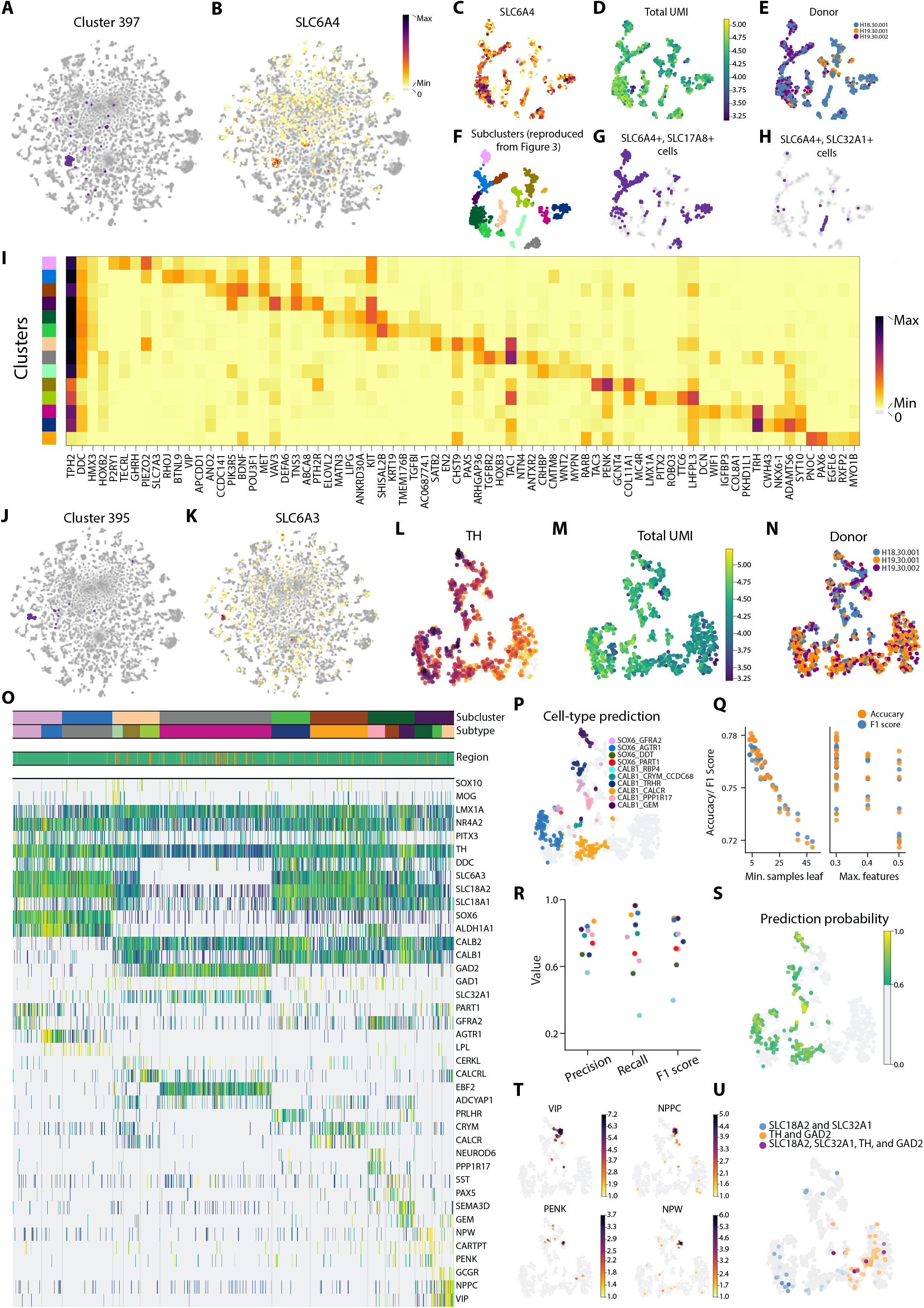
Serotonergic and dopaminergic neuronal heterogeneity within the splatter supercluster. (**A**) *Splatter* cells on the t-SNE are colored if they belong to the serotonergic cluster 397. (**B**) Cells are colored by gene expression. (**C**) Serotonergic cells are colored on the t-SNE by gene expression, legend in (B). (**D**) Cells are colored by log10 of their total UMI counts. (**E**) Cells are colored by donor. (**F**) Cells are colored by subcluster. (**G**) Cells are colored purple if they express both serotonin-reuptake transporter *SLC6A4* and glutamate transporter *SLC17A8*. (**H**) Cells are colored purple if they express both serotonin-reuptake transporter *SLC6A4* and GABA transporter *SLC32A1*. (**I**) Heatmap shows mean gene-expression levels across clusters for serotonergic markers *TPH2* and *DDC*, the rostral and caudal serotonergic markers *HMX3* and *HOXB2*, and the top five enriched genes in each cluster. Color bar on the left shows the corresponding subcluster for each row as in (F). (**J**) *Splatter* cells on the t-SNE are colored if they belong to the dopaminergic cluster 395. (**K**) Cells are colored by gene expression. (**L**) Dopaminergic cells are colored on the t-SNE by gene expression, legend in (B). (**M**) Cells are colored by log10 of their total UMI counts. (**N**) Cells are colored by donor. (**O**) A heatmap represents expression levels for select genes across dopaminergic cells. Top bar indicates the subcluster for each cell as in Fig. 3M; second bar indicates subtype as in Fig. 3P. Third bar indicates the region of origin for each cell colored as in Fig. 3N. (**P**) A random forest classifier was used to transfer labels from Kamath et. al. The classifier also reported a probability for each prediction, reflecting its confidence in the label transfer. Cells are colored if their probability was greater than 60%; other cells are grey. The colors indicate predicted label according to the legend. (**Q**) The classifier’s performance was tested at different values for two parameters: minimum number of samples permitted at a decision tree’s leaf (left) and the maximum number of features used to fit a tree (right). Performance was evaluated by both accuracy (orange) and F1 score (blue). (**R**) Precision, recall, and F1 scores are reported for each predicted label, colored according to the legend in (K). The classifier most poorly identified CALB1_RBP4 cells. (**S**) Cells on the t-SNE are colored if the classifier’s prediction probability was above 60%; other cells are grey. Color indicates probability. (**T**) Cells are colored by gene expression. (**U**) Cells are colored by the product of logged expression in multiple genes.

**Supplementary Figure 6.**
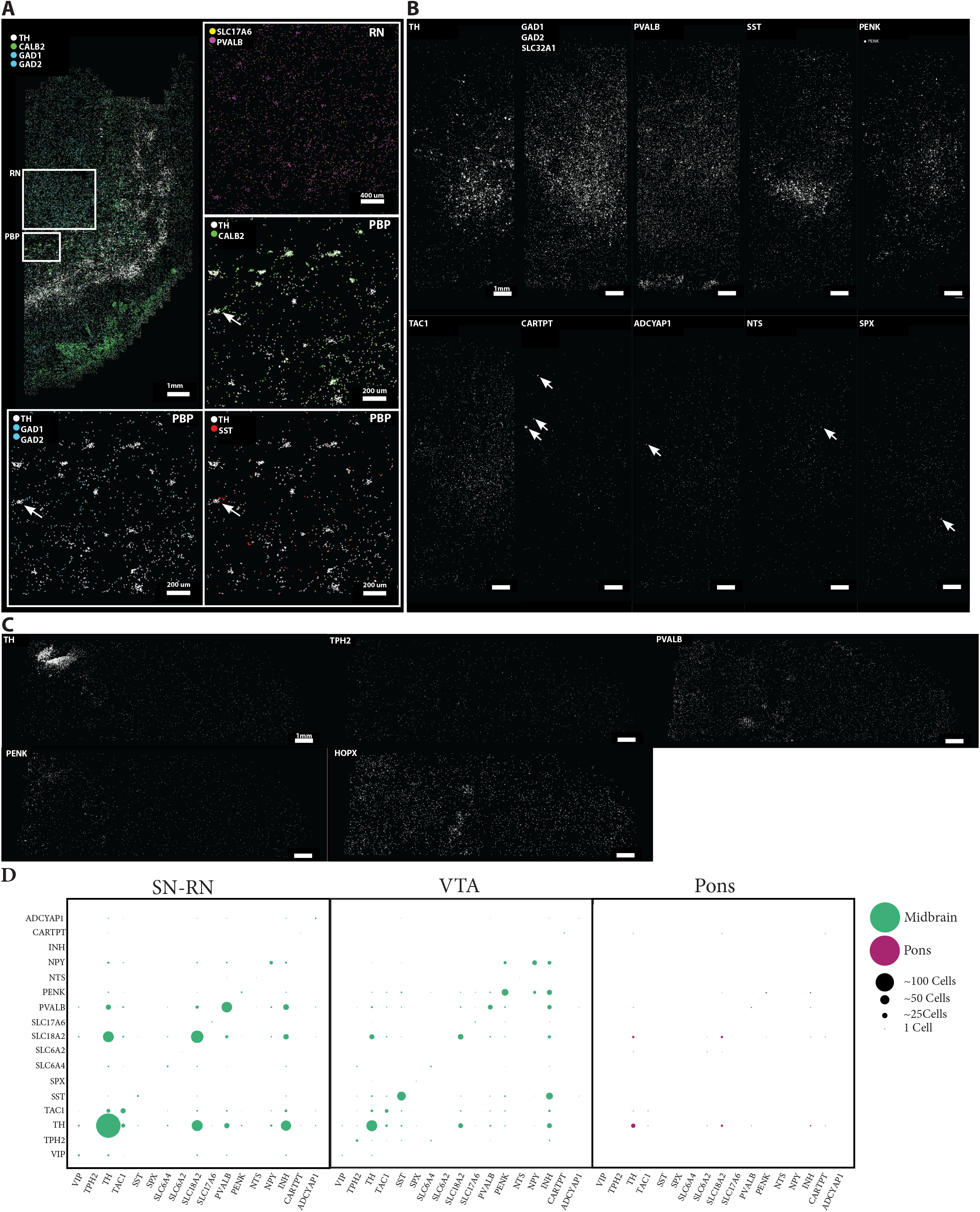
EEL-FISH profiles gene-expression distributions in the midbrain and hindbrain. (**A**) Main panel shows *TH, GAD1* or *GAD2*, and *CALB2* expression in Tissue A that includes the substantia nigra, red nucleus (RN), and parabrachial pigmented nucleus (PBP). Top right shows enlarged view of *SLC17A6* and *PVALB* co-expression in the RN. Other insets show enlarged views of cells in the PBP co-expressing *TH* with *GAD1* or *GAD2, CALB2*, and *SST*. Each dot represents a molecule. (**B**) Each dot represents a molecule of *TH, GAD1* or *GAD2* or *SLC32A1, PVALB, SST, PENK, TAC1, CARTPT, ADCYAP1, NTS*, and *SPX* in tissue B. (**C**) Each dot represents a molecule of *TH, TPH2, PVALB, PENK*, and *HOPX* in tissue section C. (**D**) For SN-RN, VTA, and pons, the size of each dot represents the number of cells that co-express two genes along the x-axis and y-axis.

**Supplementary Figure 7.**
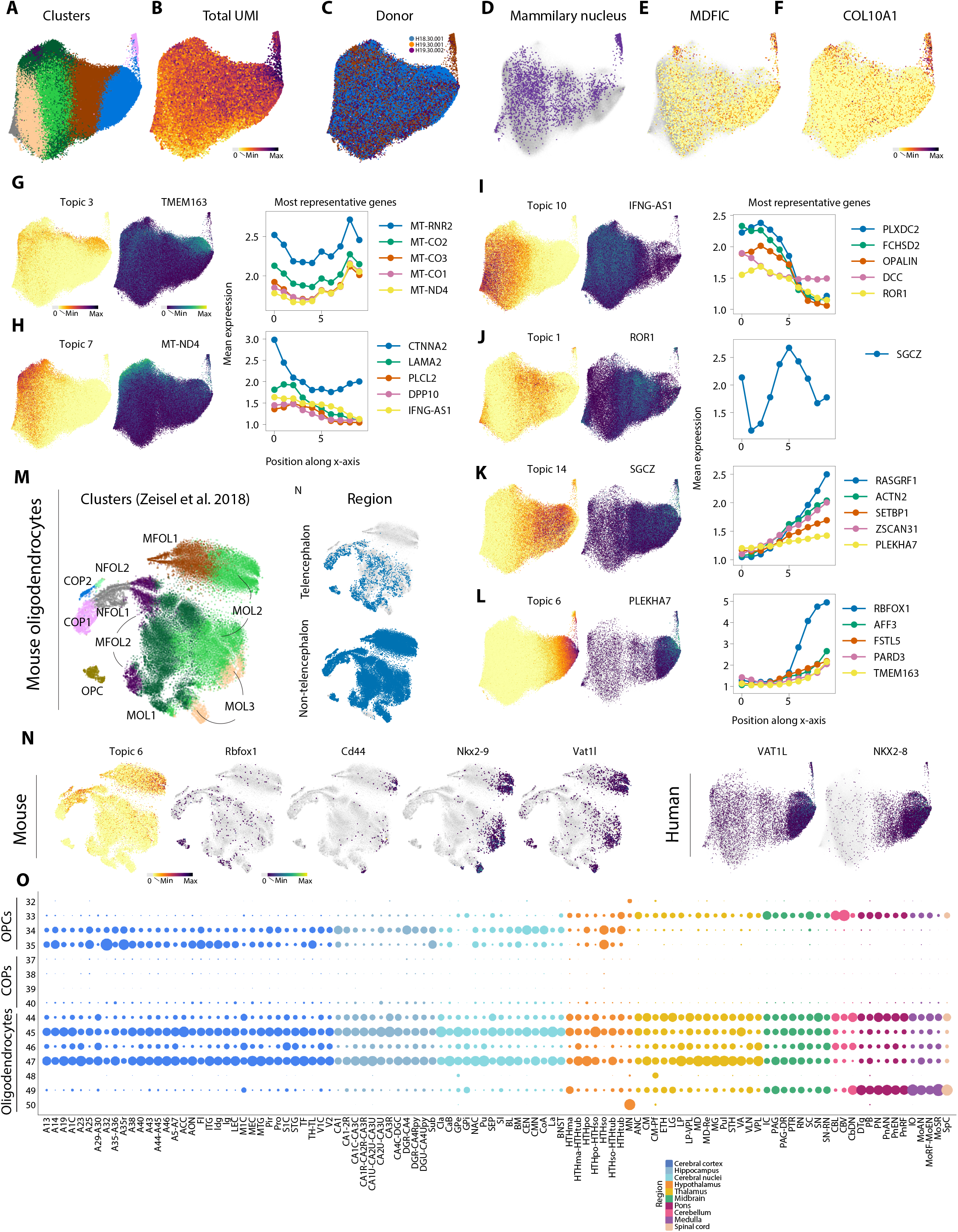
Oligodendrocyte-lineage cells differ between the telencephalon and the rest of the brain. (**A**) Oligodendrocytes on the t-SNE are colored by cluster. (**B**) Cells are colored by log10 of their total UMI counts. (**C**) Cells are colored by donor. (**D**) Cells are colored purple if they derive from the mammillary nucleus. (**E-F**) Cells are colored by gene expression. (**G-L**) Cells are colored by their scores for each gene topic (left) and their expression of a gene representative for that topic (middle). For each topic, expression values for five representative genes were binned and averaged along the t-SNE’s x-axis (right). (**M**) A t-SNE was recalculated for mouse oligodendrocytes from Zeisel et al. Cells on the t-SNE are colored by cluster. (**N**) Cells are colored blue if they derive from the telencephalon (top) or outside the telencephalon (bottom). (**N**) Mouse oligodendrocytes were scored for human Topic 6. The topic-representative gene *Rbfox1* is not expressed in mouse oligodendrocytes. Correlations were calculated between all genes and the murine Topic 6 scores in myelin-forming (MFOL) and mature (MOL) oligodendrocytes. Mouse and human cells are colored by expression of select highly correlated genes *Cd44, Nkx2-9* (*NKX2-8* in humans), and *Vat1l*. Human *CD44* expression is plotted in Fig. 5. (**O**) Dot plot represents the distributions of oligodendrocyte-lineage cells across dissections. Within each column, dot size is proportional to the percentage of cells in the indicated dissection that belong to each cluster.

**Supplementary Figure 8.**
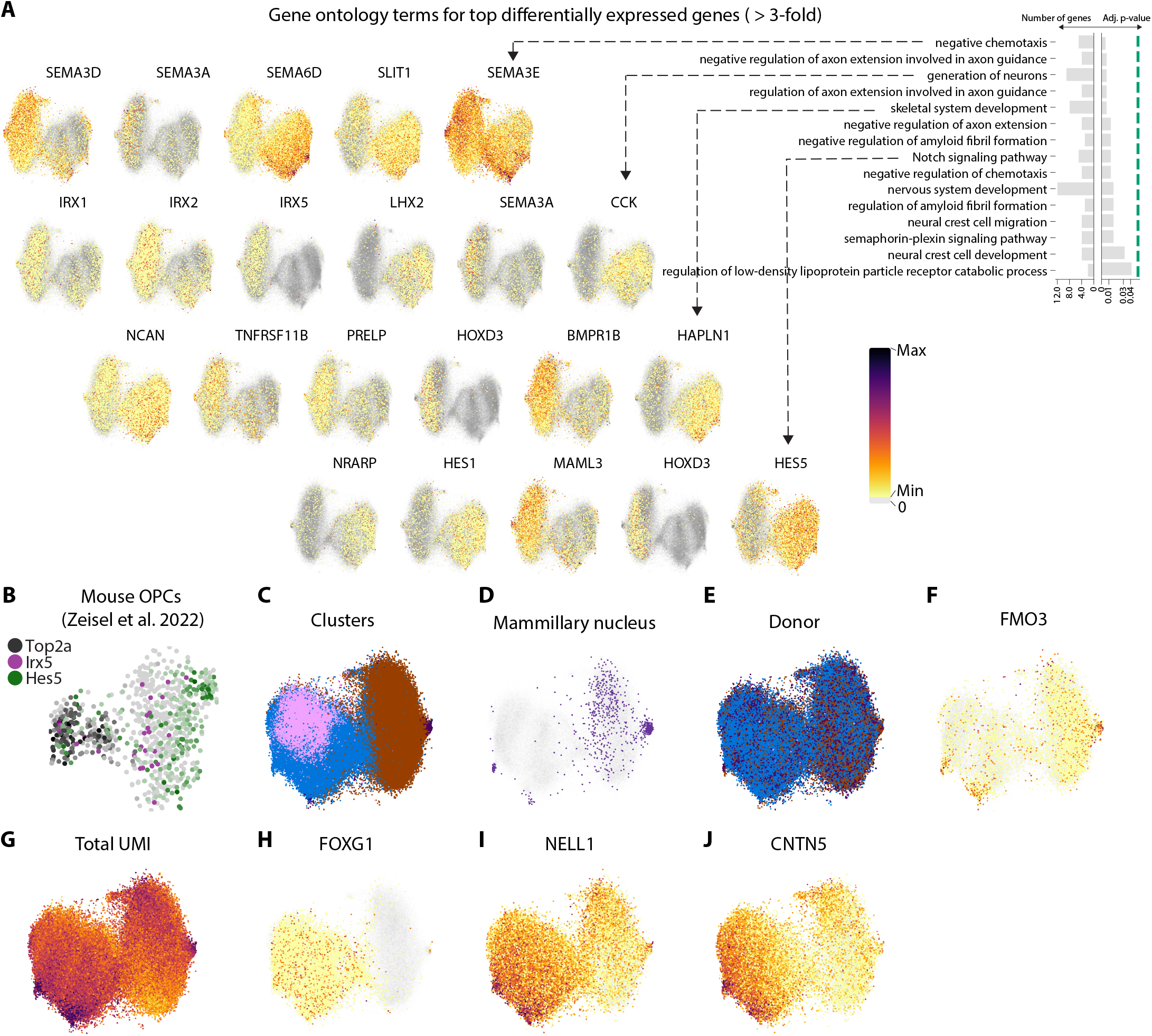
OPC heterogeneity within the oligodendrocyte lineage. (**A**) Differential expression analysis identified 155 genes with greater than three-fold enrichment in either OPC Type 1 or Type 2. On the right-hand side, bar plots show the top 15 gene-ontology terms found among the differentially expressed genes; bar lengths represents the number of genes that overlapped with a term (left) and p-value of the enrichment (right). Green line indicates 0.05. On the left-hand side, OPCs in each row are colored by their expression of genes that overlapped with the indicated gene-ontology term. (**B**) A t-SNE was calculated on OPCs from Zeisel et al. 2018. Cells are colored by expression levels of the gene they express most highly among those in the legend. Darker color indicates higher expression; gray indicates no expression. *Top2a* denotes cycling cells, and *Hes5* and *Irx5* are enriched in human Type 1 and Type2 OPCs, respectively. (**C**) Human OPCs are colored by cluster. (**D**) Cells are colored purple if they derive from the mammillary nucleus. (E) Cells are colored by donor. (**F**) Cells are colored by gene expression. (**G**) Cells are colored by log10 of their total UMI counts. (**H-J**) Cells are colored by gene expression.

**Supplementary Figure 9.**
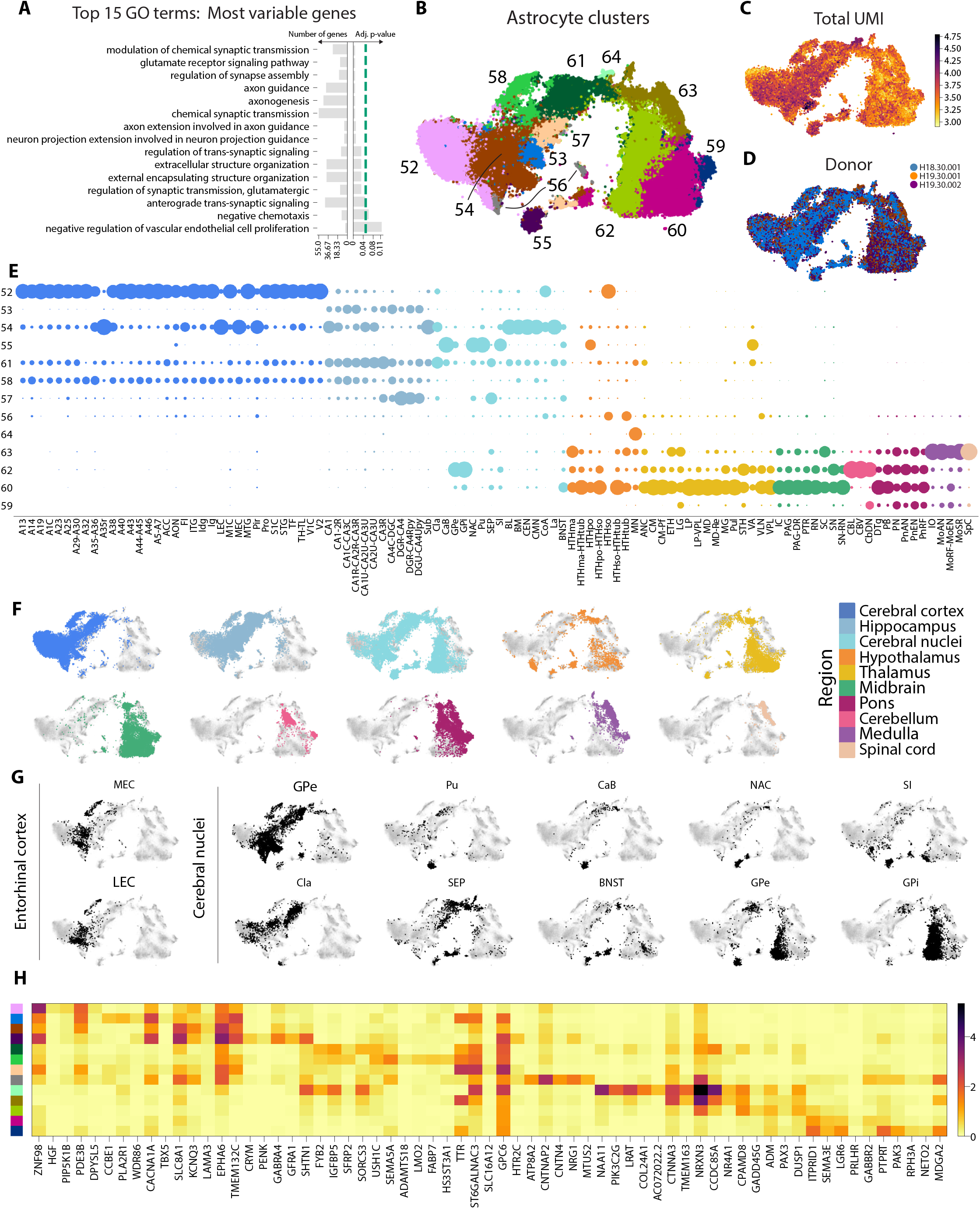
Astrocytes exhibit transcriptomic diversity across brain regions. (**A**) Bar plots show the top gene-ontology terms among the 2000 genes most variable across astrocytes. For each term, bar lengths represent the number of variable genes that overlapped with the term (left) and p-value of the enrichment (right). Green line indicates 0.05. (**B**) Astrocytes on the t-SNE are colored by cluster. (**C**) Cells are colored by log10 of their total UMI counts. (**D**) Cells are colored by donor. (**E**) Dot plot represents the distributions of astrocytes across dissections. Within each column, dot size is proportional to the percentage of cells in the indicated dissection that belong to each cluster. (**F**) Cells are colored if they derive from the region indicated by color according to the legend on the right. (**G**) Cells are colored black if they derive from the indicated dissection. (**H**) Heatmap shows mean gene-expression levels across clusters for the top five enriched genes in each cluster. Color bar on the left indicates cluster as in (B).

**Supplementary Figure 10.**
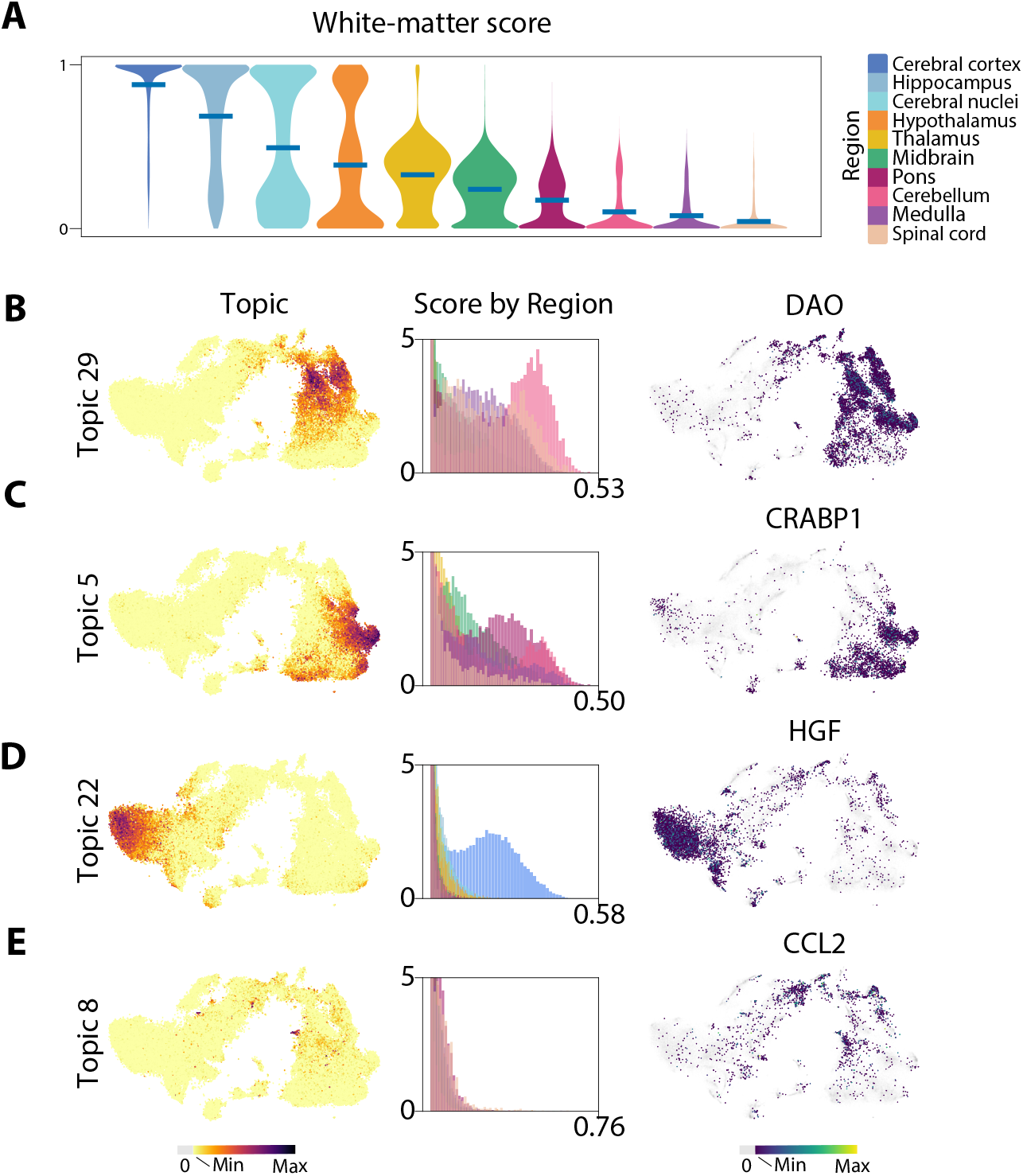
Latent Dirichlet Allocation captures astrocyte heterogeneity. (**A**) Violin plots summarize distributions of white-matter scores within each brain region. Horizontal line indicates a distribution’s mean. (**B-E**) Astrocytes are colored on the t-SNE by their scores for each gene topic (left). Scores were then split by region, and a histogram shows the score distributions for each region (middle). Cells are colored by their expression of a gene representative for that topic (right).

**Supplementary Table 1** | **Samples**. All 10x Genomics samples that were sequenced are listed with metadata in the Excel table.

**Supplementary Table 2** | **Annotations**. All clusters and their annotations are listed in the Excel table as ordered in the Figure 1A dendrogram. See methods for annotation details.

**Supplementary Table 3** | **Differential expression between Type 1 and Type 2 oligodendrocytes**. Mann-Whitney rank-test results for all genes are listed in the csv file.

**Supplementary Table 4** | **Differential expression between Type 1 and Type 2 OPCs**. Mann-Whitney rank-test results for all genes are listed in the csv file.

## Notes

https://github.com/linnarsson-lab/adult-human-brain

### Competing Interest Statement

The authors have declared no competing interest.

